# DAB2 as a biomarker and mechanistic link between lipid dysregulation and disease progression in LGMD R2

**DOI:** 10.1101/2025.09.11.675589

**Authors:** Celine Bruge, Nathalie Bourg, Emilie Pellier, Quentin Miagoux, Manon Benabides, Noella Grossi, Margot Jarrige, Helene Polveche, Valeria Agostini, Anthony Brureau, Stephane Vassilopoulos, Teresinha Evangelista, Gorka Fernández-Eulate, Tanya Stojkovic, Isabelle Richard, Xavier Nissan

## Abstract

Limb-girdle muscular dystrophy R2 (LGMD R2) is an autosomal recessive disorder caused by dysferlin deficiency, leading to progressive muscle weakness and wasting. Despite advances in understanding the mechanisms linking dysferlin loss to membrane fragility and muscle degeneration, the lack of robust clinical biomarkers has limited disease monitoring and therapeutic evaluation. Here, we identify Disabled-2 (DAB2) as a molecular and clinical biomarker for LGMD R2. Transcriptomic profiling revealed a significant upregulation of DAB2 in induced pluripotent stem cell (iPSC)-derived myotubes from patients with LGMD R2. Its expression correlated with disease severity in muscle biopsies from a cohort of 14 dysferlin-deficient individuals and in dysferlin knockout Bla/J mice, where levels increased with disease progression. Crucially, we demonstrate that DAB2 upregulation in muscle is normalized following treatment with AAV gene therapy expressing full-length dysferlin, positioning DAB2 as a dynamic biomarker for both disease monitoring and therapeutic response. Based on the role of DAB2 in lipid trafficking and the reported pathological lipid accumulation in LGMD R2, we then investigated its contribution to disease-associated lipid dysregulation. Consistent with this hypothesis, we show that high DAB2 levels paralleled lipid deposition in affected patients, iPSC-derived myotubes and mouse muscles, while siRNA-mediated DAB2 knockdown reduced lipid accumulation in LGMD R2 myotubes. Together, our findings establish DAB2 as a mechanistic link between disease severity and lipid dysregulation, and highlight its potential as a key prognostic marker, opening new avenues for precision medicine approaches in LGMD R2 and other related muscular dystrophies.

**Graphical abstract:** 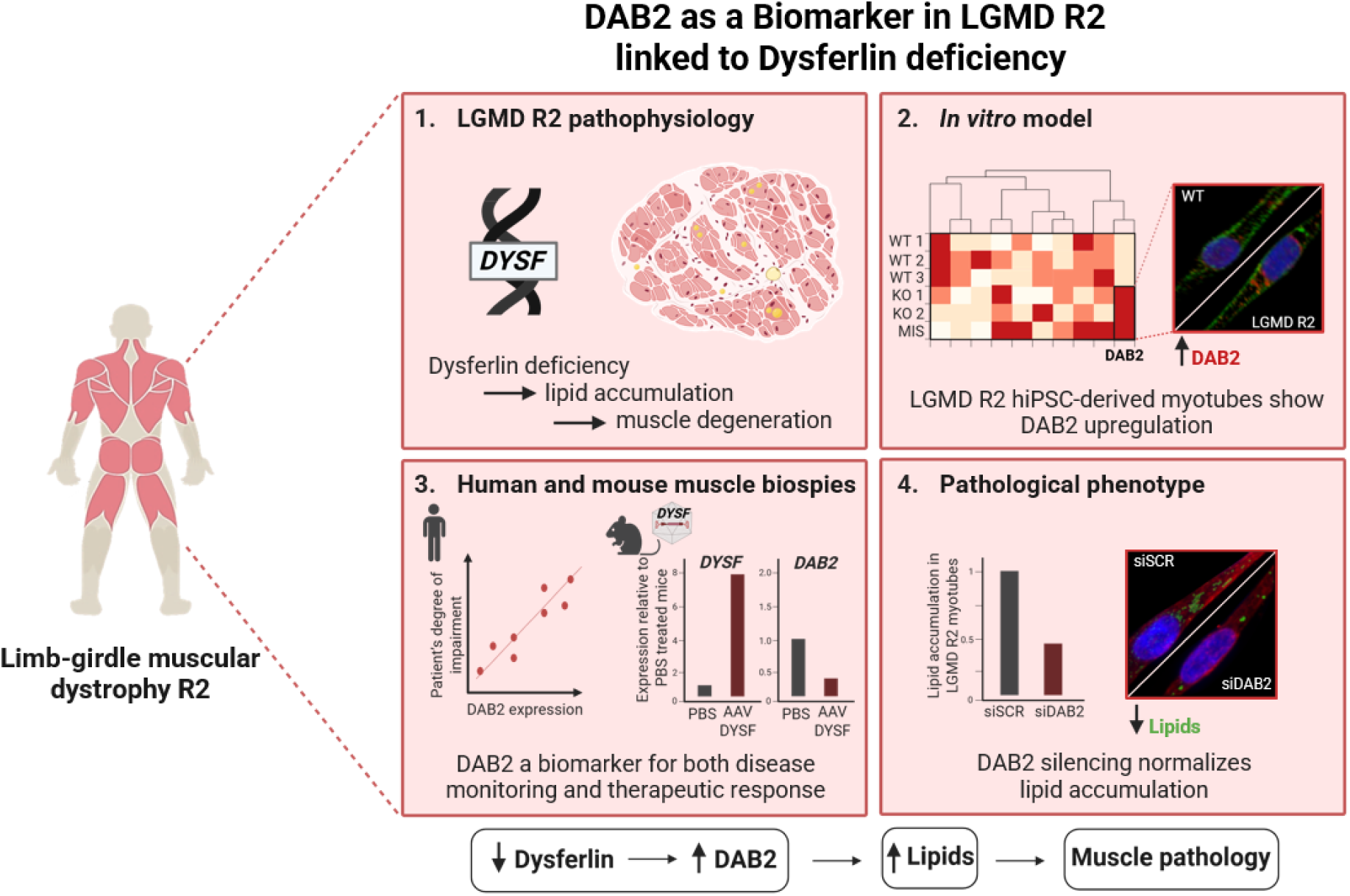

## INTRODUCTION

What triggers muscle degeneration in dysferlinopathies, and which hidden molecular pathways drive its progression? Here, we sought to investigate these underlying mechanisms at the molecular level. Dysferlinopathies comprise a genetically heterogeneous group of muscular dystrophies. Among them, limb girdle muscular dystrophy R2/2B (LGMD R2/2B) is characterized by progressive weakness and wasting of the pelvic and scapular girdle muscles. With an estimated prevalence of 5.9–7.4 individuals per million worldwide (1), LGMD R2 is caused by autosomal recessive mutations in the *DYSF* gene (2, 3), which encodes dysferlin, a 230-kDa transmembrane protein highly expressed in striated muscle and predominantly localized at the sarcolemma. While several approaches have been described to partially restore muscle strength or motor function in preclinical models (4–16), no curative therapy is currently available, highlighting the need to better understand the molecular mechanisms underlying dysferlin deficiency.

Historically, studies aiming to elucidate the pathological mechanisms underlying dysferlinopathies have focused primarily on the role of dysferlin in plasma membrane repair (PMR) of the injured sarcolemma; a calcium-dependent process critical for maintaining myofiber integrity and function (17). While defects in PMR are observed in muscle fibers from dysferlin-deficient mice (18) and patient myocytes (19), studies also reported that its restoration alone (6) is insufficient to fully prevent or reverse dystrophic phenotypes (7), indicating that additional mechanisms contribute to disease progression. In agreement with these observations, several studies have then highlighted an expanded role of dysferlin through its involvement in t-tubule formation and maintenance (20–24), intracellular vesicular trafficking (25), and calcium homeostasis (20–22, 24, 26–28).

More recently, evidence from patient muscle biopsies and dysferlin-deficient mouse models indicated that progressive skeletal muscle remodeling is preceded by intramyofiber lipid accumulation, followed by adipocyte infiltration between fibers (29–32) proposing lipid metabolism as an early driver of LGMD R2 pathogenesis (33). This hypothesis was further supported by several studies (34–37) indicating that cholesterol homeostasis and lipid accumulation are not merely a secondary consequence of muscle degeneration, but an early, intrinsic, and potentially used as a therapeutical target in LGMD R2.

To dissect how early molecular defects drive progressive muscle damage, robust models that faithfully recapitulate human dysferlinopathy are essential. In the past years, a large number of studies, including from our group, have demonstrated the relevance of human induced pluripotent stem cells (hiPSC) for the identification of new molecular mechanisms and pathological phenotypes of muscular diseases (38–45). Building on this approach, we generated skeletal muscle cells (skMC) from three LGMD R2 patient-derived hiPSC to investigate the cellular and molecular processes involved in this pathology. Our approach started from the identification of Disabled-2 (*DAB2*) as consistently upregulated in LGMD R2 hiPSC-derived myotubes using a comparative transcriptomic analysis. Measures of DAB2 expression in muscle biopsies from a cohort of LGMD R2 patients and dysferlin knockout Bla/J mice confirmed its overexpression in multiple models and further revealed that it correlates with lipid accumulation, disease severity, and progression. In this study we also showed that dysferlin restoration normalized DAB2 levels, demonstrating that its expression reflects both disease burden and therapeutic response. To directly link DAB2 with lipid accumulation, we finally performed loss of function experiments to explore their mechanistic link. Collectively, these findings position DAB2 as both a clinically relevant biomarker and a potential regulator of lipid metabolism in LGMD R2, providing new insights into disease mechanisms and identifying a candidate therapeutic target for gene therapy or pharmacological interventions.

## RESULTS

### Characterization of LGMD R2 hiPSC-derived skeletal muscle cells (skMC)

We previously reported dysferlin expression in healthy hiPSC-derived skMC (44). To investigate the molecular mechanisms underlying LGMD R2, we differentiated three dysferlin- mutant hiPSC lines using the same standardized protocol (45). Two lines carry nonsense mutations (LGMD R2 KO_1 and KO_2), resulting in complete loss of dysferlin isoforms, whereas a third line harbors a missense mutation (LGMD R2 MIS) producing an aggregated, misfolded dysferlin protein, as previously reported (46). Pluripotency of all LGMD R2 hiPSC lines was confirmed by measuring SSEA4 and TRA-1-81 expression using flow cytometry (Figure S1). Both LGMD R2 and control hiPSC were then differentiated using the established protocol (Figure 1A). Quantitative PCR showed no significant differences in *OCT4* (Figure S2A) or *NANOG* (Figure S2B) pluripotency markers expression between control and LGMD R2 hiPSC lines (day 0). Throughout differentiation, the downregulation of pluripotency markers (Figures S2A and S2B) and induction of myogenic markers (*MYOD*, *DESMIN*, *MYOG*, *TTN;* Figures S2C-S2G) occurred with similar kinetics in both LGMD R2 and control cells, indicating that dysferlin deficiency does not impair early or late stage of myogenic differentiation. Additionally, immunostaining confirmed the ability of LGMD R2 hiPSC to form a well-organized network of striated myotubes positive for titin (Figure S2H) and myosin heavy chain (Figure S2I), similarly to unaffected control cells. Immunoblotting demonstrated complete absence of dysferlin in myoblasts and myotubes derived from the LGMD R2 nonsense lines and a marked reduction in the missense line compared with controls (Figures 1B-1C). These findings were confirmed by immunostaining, which showed absence of dysferlin in KO myoblasts (Figure 1D) and myotubes (Figure 1E), and perinuclear accumulation in MIS myotubes (Figure 1E).

**Figure 1.**
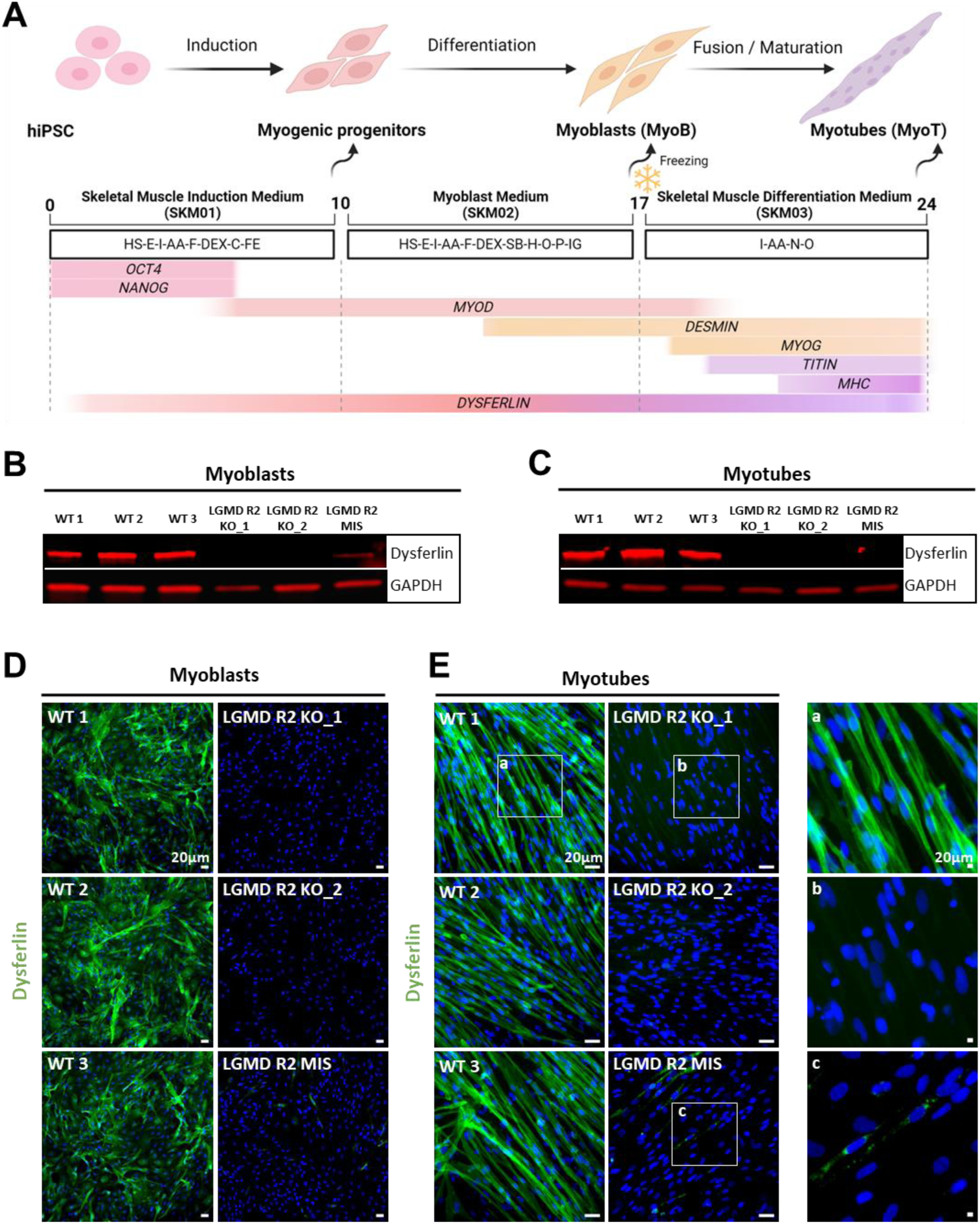
Phenotypic features of LGMD R2 hiPSC-derived skeletal muscle cells reveal dysferlin deficiency. (**A**) Schematic representation of the skeletal myogenic differentiation protocol. Arrows indicate key stages where specific phenotypes are observed. Growth factors and cytokines used at each differentiation stage are specified, along with the progression of selected gene expression. Schematic created with BioRender. (**B-C**) Immunoblot analysis of dysferlin expression in LGMD R2 and control hiPSC-derived myoblasts (**B**) and myotubes (**C**). GAPDH is used as a loading control. (**D-E**) Immunostaining of dysferlin (green) in LGMD R2 and control myoblasts (**D**) and myotubes (**E**). Nuclei were counterstained with Hoechst (blue). Insets (a–c) show magnified views of dysferlin staining in myotubes. Scale bar = 20µm. Abbreviations: AA, ascorbic acid; C, CHIR99021; Dex, dexamethasone; E, epidermal growth factor; F, basic fibroblast growth factor; Fe, fetuin; H, hepatocyte growth factor; HS, horse serum; I, insulin; IG, insulin-like growth factor; N, necrosulfonamide; O, oncostatin; P, platelet-derived growth factor; SB, SB431542; hiPSC, human induced pluripotent stem cells.

### Transcriptomic profiling reveals gene signatures and disrupted pathways in LGMD R2 myotubes

To investigate global transcriptional changes, we performed RNA sequencing on LGMD R2 and control hiPSC-derived myotubes. Principal component analysis (PCA) and heatmap analysis revealed distinct clustering of LGMD R2 and control samples (Figures S3A and 2A). Differential expression analysis identified 52 differentially expressed genes (DEGs), including 13 downregulated and 39 upregulated genes in LGMD R2 myotubes (Figure 2B). Quantitative PCR analysis validated 20 of 23 selected DEGs, including *DYSF* itself (Figures S3B-S3C). Finally, enrichment analysis using Enrichr revealed that downregulated genes were associated with fibrosis-related pathways (e.g., TGF-beta signaling) and lipid metabolism (Figure S3D), while upregulated genes were linked to extracellular matrix (ECM) remodeling (Figure S3E). Gene Ontology (GO) analysis further highlighted relevant biological processes, such as ‘regulation of intracellular transport’ and ‘transmembrane receptor protein’, with a gene network centered on *DAB2* (Figures 2C-2D).

**Figure 2.**
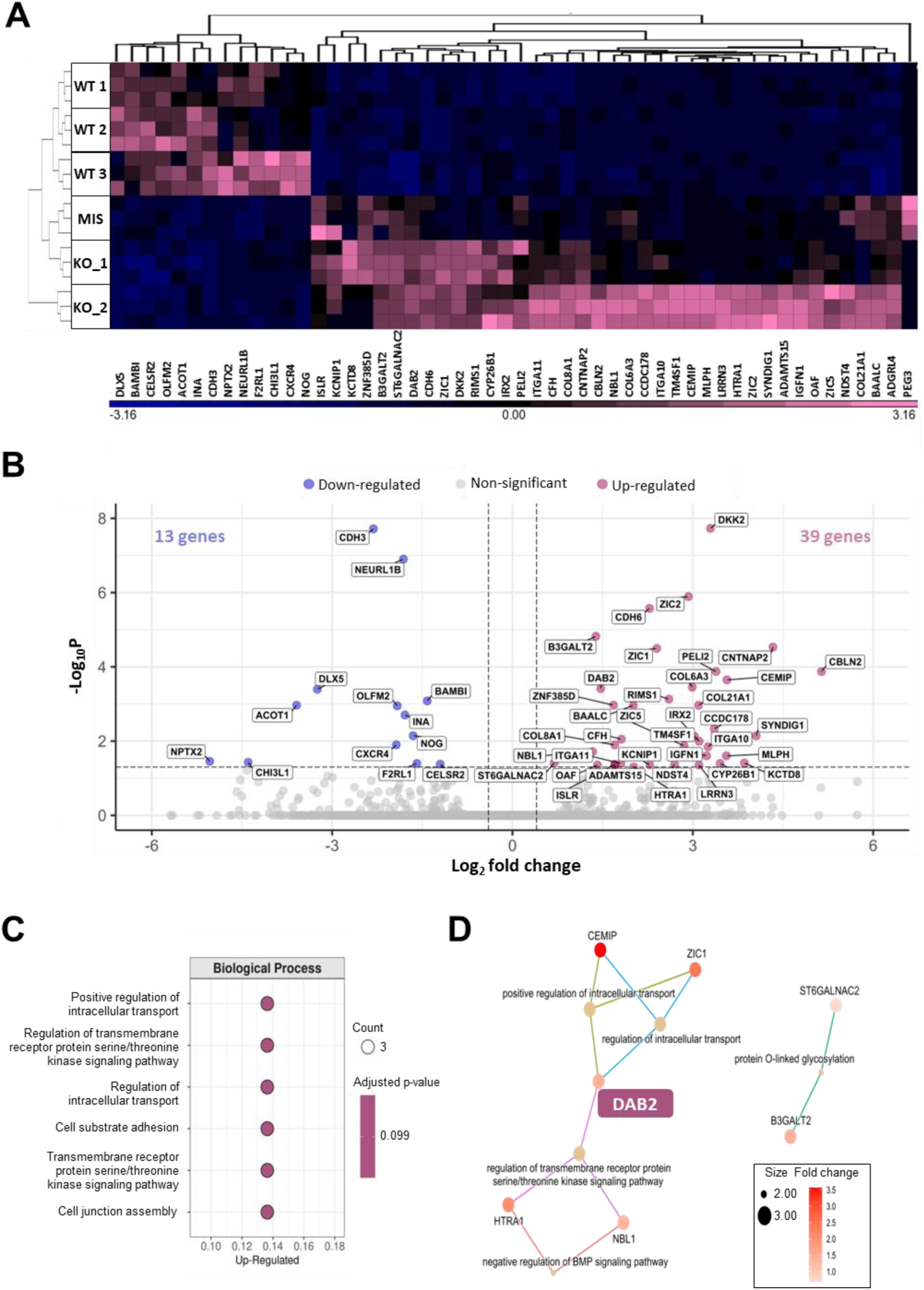
Transcriptomic profiling reveals altered gene expression signatures in LGMD R2 hiPSC-derived myotubes. (**A**) Hierarchical clustering of differentially expressed genes detected in LGMD R2 myotubes compared with controls. Gene expression is color-coded from blue (downregulated) to pink (upregulated). (**B**) Volcano plot representation of differential gene expression analysis between myotubes derived from three control and three LGMD R2 hiPSC lines. Downregulated (blue) and upregulated (pink) genes in LGMD R2 myotubes are highlighted. Vertical dashed lines indicate |log2Foldchange|threshold ≥ 0.4, and the horizontal dashed line represents a false discovery rate (FDR) ≤ 0.05. (**C**-**D**) Enriched gene ontology (GO) terms obtained by over- representation analysis (ORA) of upregulated genes in LGMD R2 myotubes compared with controls. Each pathway has an adjusted p-value of 0.0995. (**C**) Dot plots show the six GO terms shared by at least three genes. (**D**) Cnetplot of the five biological processes most significantly shared among GO terms. Bubble size represents the number of enriched genes. DAB2 is highlighted in pink.

### DAB2 as a marker of LGMD R2 pathophysiology

DAB2 overexpression in dysferlin-deficient myotubes was first confirmed at the transcript level by qPCR (Figure 3A) and validated at the protein level by western blotting (Figure 3B). Confocal imaging further demonstrated increased cytoplasmic accumulation of DAB2, as evidenced by a higher number of DAB2-positive puncta in LGMD R2 myotubes compared with control (Figure 3C, Movies S1-S2). To assess the relevance of these findings in human disease, we analyzed DAB2 expression in skeletal muscle biopsies from fourteen genetically confirmed dysferlin-deficient patients and two unaffected controls (Table S1). Dysferlin pathogenic variants were distributed across the protein without clustering at specific hotspots (Figure 3D). In most patient samples, DAB2 mRNA was significantly elevated in LGMD R2 patient biopsies compared with controls (Figure 3E). These findings were further supported by analysis of a previously published RNA sequencing dataset from ten LGMD R2 patients and thirteen unaffected controls (47, 48), where we analyzed and confirmed the increase DAB2 expression in most patient muscle biopsies (Figure S4). Importantly, our analysis confirmed that DAB2 expression did not correlate with patient gender (Figure S5A), clinical subtype of dysferlinopathy (Figure S5B), or serum creatine kinase (CK) levels (Figure S5C). Instead, DAB2 levels closely reflected the severity of muscle pathology. Patients with more advanced disease stages, indicated by higher Walton scores (Figure 3F) and pronounced morphological muscle alterations (Figure S5D), displayed the highest DAB2 expression. Protein-level analysis confirmed these findings: DAB2 was markedly increased in the muscle biopsy from patient 2, who exhibited clear dystrophic features, while no modulation was observed in patient 6, whose muscle appeared morphologically normal and comparable to a healthy control (Figure 3G). Collectively, these data support the use of DAB2 as a marker of dysferlin deficiency and disease progression.

**Figure 3.**
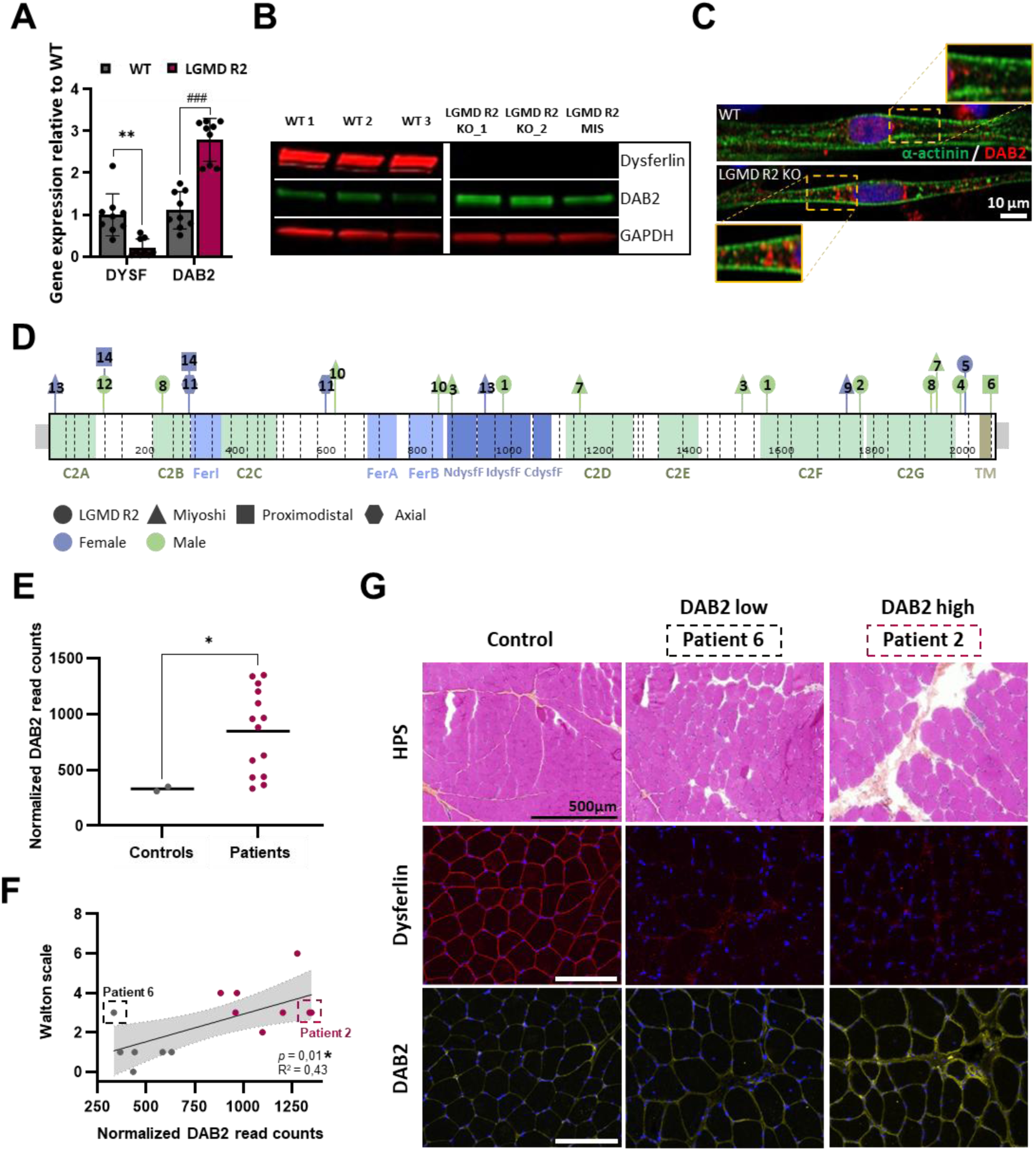
Dysregulation of Disabled-2 (*DAB2*) in human dysferlin-deficient myotubes and muscle biopsies. (**A-C**) Analysis of DAB2 in LGMD R2 hiPSC-derived myotubes. (**A**) Quantitative PCR analysis of *DYSF* and *DAB2* expression in myotubes from three control (grey) and three LGMD R2 (pink) hiPSC lines. Data represent mean ± SD of three independent differentiations per line (n = 9), normalized to the mean of control myotubes. **p ≤ 0.001 (unpaired t test with Welch’s correction); ###p ≤ 0.001 (unpaired t test). (**B**) Immunoblots of dysferlin (red) and DAB2 (green) in control and LGMD R2 myotubes. GAPDH served as loading control. (**C**) Immunostaining of DAB2 (red) and α-actinin (green) in control (top) and LGMD R2 (bottom) myotubes. Nuclei were counterstained with Hoechst (blue). Insets show magnified regions. Scale bar = 10 µm. (**D-G**) DAB2 expression in muscle biopsies from 14 dysferlin-deficient patients. (**D**) Map of patient mutations in *DYSF* gene (exons separated by dotted lines). Protein domains are indicated. Each patient is numbered; clinical phenotype and sex are indicated by shape and color (see Supplementary Table 2). (**E**) DAB2 mRNA levels from QuantSeq (Illumina) in control (grey) and patient (pink) muscle biopsies. Each dot represents a biopsy; bars indicate mean. *p ≤ 0.05 (Mann–Whitney test). (**F**) Correlation between DAB2 expression and patient involvement (mildly affected 0 ≤ Walton scale ≤ 10 severely affected). Dots represent biopsies, color-coded from grey (low DAB2) to pink (high DAB2). Patients with the lowest (patient 6) and highest (patient 2) DAB2 expression are highlighted. *p ≤ 0.05 (Pearson correlation). (**G**) HPS-stained deltoid sections from patients 6 and 2 with immunostaining for dysferlin (red) and DAB2 (yellow). Scale bar = 500 µm. Abbreviations: HPS, hematoxylin phloxine saffron. Protein domains: CdysfF, dysferlin domain C-terminal region; Fer, Ferlin domain; IdysfF, inner dysferlin domain; NdysfF, dysferlin domain N-terminal region; TM, transmembrane domain.

### DAB2 expression inversely correlates with dysferlin content *in vivo*

We next assessed Dab2 expression in dysferlin-deficient Bla/J mouse model (4). Homozygous Bla/J mice develop a progressive muscular dystrophy and recapitulate the main histopathological features observed in patients, including inflammation, fiber degeneration, and centrally nucleated fibers. Dystrophic changes are detectable at the age of 8 weeks and worsen by the age of 4 months, with widespread muscle impairment by 8 months. To investigate Dab2 dynamics at early disease stages, we analyzed psoas and gluteus muscles at 3 and 6 months of age. Tibialis anterior, which remains preserved in this model was used as a control. At 3 months, histological analysis revealed that the psoas was the most affected muscle, with few centrally nucleated fibers and inflammatory areas (Figure 4A). By 6 months, both psoas and gluteus, but not tibialis anterior, showed significant morphological alterations (Figures 4A and S6). qPCR analysis of Dab2 confirmed the correlation with disease progression as shown with the significant increase in Dab2 mRNA in the psoas at 3 months, with no changes in gluteus or tibialis anterior (Figure 4B) and its upregulation at 6 months in both psoas and gluteus, but not in tibialis anterior (Figure 4C). These results were confirmed at the protein level by immunoblotting, which showed overexpression of Dab2 isoforms in psoas and gluteus muscles of 6-month-old Bla/J mice compared to controls (Figure 4D). To evaluate the use of Dab2 expression as a therapeutic biomarker, we next analyzed its expression in dysferlin-deficient pre-symptomatic mice treated with a dual AAV vector delivering full-length dysferlin (8) (Figure S7A). Histological analysis showed that dysferlin rescue prevented dystrophic features in psoas and gluteus at both 1- and 6-months post-injection, compared with saline-treated mice (Figures 4E and S7B). Correspondingly, qPCR revealed that dysferlin rescue (Figures 4F and S7C) was associated with a significant normalization of Dab2 mRNA levels (Figures 4G and S7D) in 1 and 6 months-treated mice. These results suggest that Dab2 can be used to monitor disease progression and therapeutic efficacy *in vivo*.

**Figure 4.**
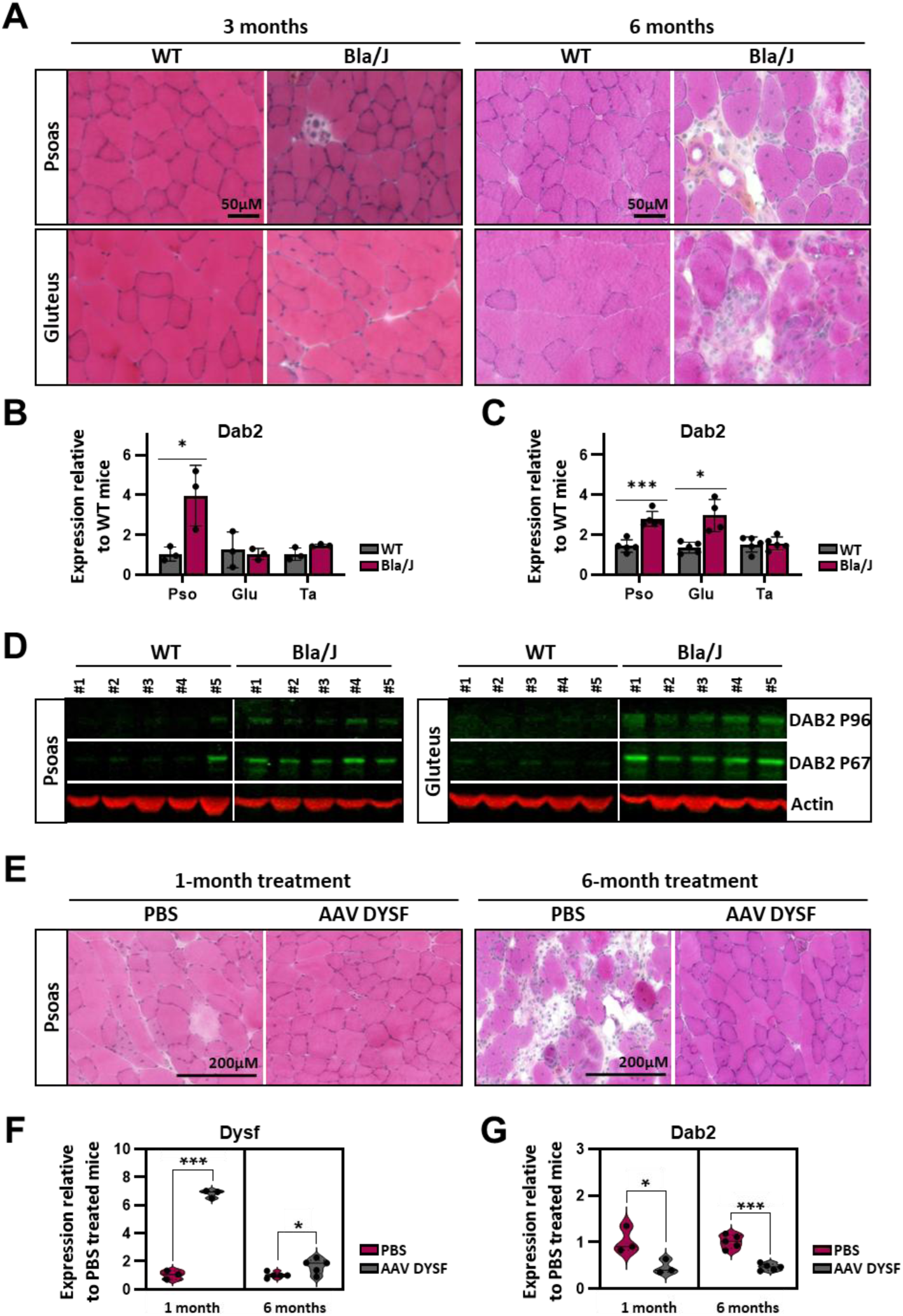
DAB2 is upregulated in severely affected muscles of dysferlin-deficient mice and restored upon dysferlin gene therapy. (**A-D**) Analysis of Dab2 expression in dysferlin-deficient mice model (Bla/J). (**A**) Histological comparison of psoas and gluteus muscles from three-month-old (left) and six-month-old (right) control and Bla/J mice using HPS staining. Scale bar = 50 µm. Associated quantification of Dab2 mRNA by qPCR in psoas, gluteus, and tibialis anterior muscles of control (grey) and Bla/J (pink) mice at three (**B**) and six months (**C**). Expression levels are normalized to control mice. Data represent mean ± SD (n = 3–5 per group). *p ≤ 0.05, ***p ≤ 0.001 (unpaired t test with Welch’s correction). (**D**) Immunoblots of murine Dab2 isoforms (green) in psoas (left) and gluteus (right) muscles from six-month-old control and Bla/J mice. Actin is a loading control. (**E- G**) Dab2 expression following AAV-mediated dysferlin gene therapy in Bla/J mice. (**E**) Representative HPS- stained psoas sections from one-month-old Bla/J mice after one month (left) or six months (right) of PBS or AAV dysferlin treatment. Scale bar = 200 µm. Associated measure of *Dysf* (**F**) and *Dab2* (**G**) mRNA by qPCR in psoas muscles of PBS-treated (pink) or AAV-dysferlin-treated (grey) Bla/J mice. Expression levels are normalized to PBS-treated mice. Data represent mean ± SD (n = 5 per group). *p ≤ 0.05, **p ≤ 0.01, ***p ≤ 0.001 (unpaired t test with Welch’s correction). Abbreviations: Pso, psoas muscle; Glu, gluteus muscle; Ta, tibialis anterior muscle.

### DAB2 expression is associated with lipid accumulation in dysferlin-deficient muscle and myotubes

Based on the reported defective lipid metabolism in LGMD R2 and the role of DAB2 in low-density lipoprotein receptor (LDLR) endocytosis and cholesterol-rich LDL uptake (49, 50), we then assessed the relation between dysferlin deficiency, lipid accumulation and DAB2 expression. To do so, we first temporally and spatially characterized these parameters in Bla/J mice. Histological analysis of Bla/J mice confirmed that lipid deposition occurred predominantly in affected muscles, with strong Oil Red O staining in psoas and gluteus, but not tibialis anterior, at 6 months (Figure S8). Dysferlin gene therapy markedly reduced lipid accumulation in the psoas (Figure 5A) and gluteus (Figure S9) after 6 months of treatment compared with PBS-treated controls, consistent with previous findings. These observations were then confirmed in low, high and very high DAB2 expressing patient muscles. Measure of lipid accumulation by Oil Red O staining revealed that patients exhibiting high or very high DAB2 content showed an increased lipid deposition. Inversely, patients with low DAB2 levels and healthy controls displayed minimal lipid accumulation (Figure 5B). These findings were finally confirmed *in vitro* by analyzing lipid uptake in hiPSC-derived myotubes treated with fatty acids. Analysis of LGMDR2 and control cell lines revealed that LGMD R2 myotubes exhibited significantly greater lipid accumulation than controls, as shown by immunostaining (Figures 5C and S10A–E) as well as by the quantification of lipid droplet volume (Figure 5D) and number (Figure S10F) in myotubes. We finally performed functional analyses to assess the role of DAB2 in lipid accumulation. To do so, siRNA-mediated knockdown of DAB2 was performed in hiPSC-derived myotubes (Figures S10G). Lipid measurements in siRNA-treated LGMD R2 cells revealed reduced lipid deposition (Figures 5E and S10H). Quantitative analyses confirmed significant decreases in lipid droplet volume (Figures 5F and S10J) upon DAB2 silencing, consistent with the observed downregulation of DAB2 protein levels (Figures S10H-I). Together, these results support a functional link between DAB2 expression and pathological lipid accumulation in various dysferlin-deficient models.

**Figure 5.**
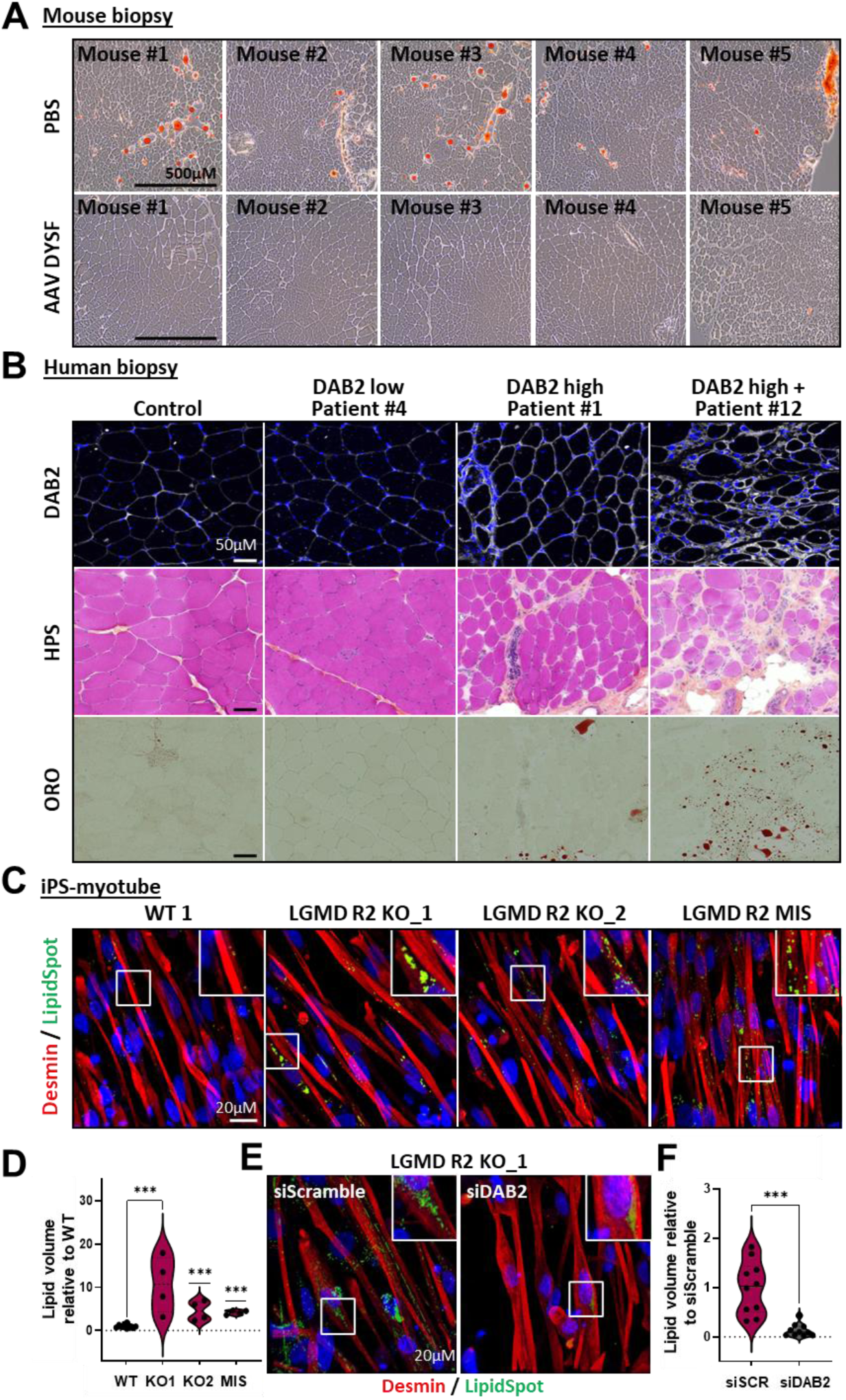
DAB2 expression is associated with lipid accumulation in dysferlin-deficient muscle and myotubes. (**A**) Oil Red O-stained sections of psoas muscles from one-month-old Bla/J mice treated for six months with PBS or AAV-dysferlin (AAV DYSF). Scale bar = 500µM. (**B**) Deltoid biopsies from a healthy control and dysferlin-deficient patients with low (patient 4), high (patient 1), or very high (patient 12) DAB2 expression, stained for DAB2, HPS, and ORO. Scale bar = 50µm. (**C-F**) Lipid accumulation in hiPSC-derived myotubes supplemented with fatty acids. (**C**) Immunostaining of lipid droplets (green) in myotubes (red) from one control and three LGMD R2. Nuclei were counterstained with Hoechst (blue). White boxes indicate magnified regions. Scale bar = 20 µm. (**D**) Associated quantification of lipid volume in control and LGMD R2 myotubes. Data represent mean ± SD of one representative experiment from *n*=3 independent experiments. ***p ≤ 0.001 (one-way ANOVA with Dunnett’s multiple comparisons test after transformation). (**E**) Immunostaining of lipid droplets (green) in LGMD R2 KO_1 myotubes transfected with siScramble (left) or siDAB2 (right). White boxes indicate magnified regions. Scale bar = 20 µm. (**F**) Associated quantification of lipid volume in LGMD R2 KO_1 myotubes after treatment with siScramble (pink) or siDAB2 (grey). Data represent mean ± SD of one representative experiment from *n*=3 independent experiments. ***p ≤ 0.001 (unpaired t test with Welch’s correction). Abbreviations: HPS, hematoxylin phloxine saffron; ORO, Oil Red O; siSCR, siScramble.

## DISCUSSION

In this study we investigated the molecular mechanisms associated with dysferlin deficiency and identified the upregulation of DAB2 in cellular and animal models of the disease and in patient muscle biopsies. We also revealed the correlation of DAB2 expression with the disease severity and the robustness of this biomarker to measure the efficacy of therapeutic interventions.

While LGMD R2 is caused by a genetic mutation, predicting functional decline of the patients or therapeutic efficacy is a challenge because of the slow and often variable progression of the disease. In the past decades, increasing efforts have been made in identifying clinical non- invasive biomarkers for muscular dystrophies and in LGMD R2 specifically. So far, the most common biological marker in muscular dystrophies is the serum CK level, an energy metabolism enzyme that leaks from damaged muscles (51). In patients with dysferlinopathy, CK levels in the blood are usually elevated (52, 53) and this elevation persists throughout the course of the disease. However, CK levels present variations due to several external conditions (physical activity, muscle injury, toxic agents, age…). Thus, although serum CK measurement is a useful diagnostic biomarker for muscular dystrophies, it is not appropriate for the assessment of disease progression. More recently, three proteins were reported to be significantly elevated in the serum of LGMD R2 patients and to correlate with the time needed to walk 10 meters in patients (54). The identified proteins include skeletal troponin I (sTnI), myosin light chain (MYL3) and fatty acid binding protein 3 (FABP3), that were also identified to be elevated in patients with Duchenne muscular dystrophy (DMD). The creatine/creatinine metabolite ratio identified in DMD was also reported to be elevated in serum of LGMD R2 patients (55). Further investigations are needed, however, to determine whether one or several (i.e. a group) of these circulating biomarkers is particularly useful for monitoring the clinical outcomes of patients with dysferlinopathy. Indeed, as recently demonstrated with two muscle growth factors, follistatin and myostatin, two biomarkers whose level is consistent with the patient’s motor function and muscle condition at initial assessment is not systematically correlated with disease progression (56). Here, we report promising results identifying DAB2 as a potential biomarker of disease severity in dysferlinopathies. Our findings revealed that DAB2 overexpression in mice muscles was exacerbated in the most affected muscles and correlated with the disease progression. In the same way, our results show that DAB2 expression levels in patient biopsies correlated with the patient’s Walton scale, paving the way for the use of DAB2 as a molecular biomarker and prognostic tool. Our study also demonstrated that DAB2 transcript level is rescued following dysferlin re-expression in KO-DYSF mice. Beyond the use of DAB2 as a prognostic tool, these results suggest its use as a pharmacodynamic biomarker to assess the benefits of drug candidates or therapeutic approaches. Indeed, with recent progress in pharmacotherapy or gene therapy for muscular dystrophies, there is a growing need for minimally invasive biomarkers that can be used to assess and monitor the efficacy of new treatments in clinical trials. Current methods include functional evaluation scales to measure patient status, measurement of the level of fatty infiltration by MRI and quantification of serum proteins (57). While promising international studies are underway to identify the most relevant outcome measures with a view to conducting clinical trials in dysferlinopathies (53, 57), so far there is only one protein that has been identified as a potential biomarker for monitoring the outcome of therapeutic interventions i.e., the myofibrillar structural protein myomesin-3 (MYOM3)(58).

Together our findings report a significant inverse correlation between elevated DAB2 expression and low dysferlin content. However, despite these observations, correlation does not inherently imply causation and determining a functional role of DAB2 in LGMD R2 could open new opportunities for the development of future therapies. In the past decade, DAB2 has been described in the literature as a bona-fide clathrin adaptor molecule involved in clathrin- mediated endocytosis (CME) (59). CME generates small (60–120 nm) membrane vesicles that transport various cargo molecules from the plasma membrane to endosomes. Importantly, DAB2 has been shown to act as an adaptor molecule for internalization of only a subset of receptors endocytosed by CME (60) and notably involved in LDLR internalization and cholesterol-rich LDL uptake (49, 50). The uptake of LDL is highly responsive to even small changes of cholesterol concentrations and deprivation of cholesterol induces SREBP transcription factor activation, which in turn regulates LDLR expression (61). Interestingly, in the past years, several studies have reported the significant accumulation of a broad range of lipids in pre-symptomatic dysferlin-deficient mice (31) and a deleterious effect of non HDL-associated cholesterol in dysferlin-deficient muscles (36, 62). Additionally, recent comprehensive lipidomic analyses revealed widespread alterations in both storage and membrane-associated lipids, including neutral lipids, phospholipids, and sphingolipids, coupled with reduced fatty acid oxidation and a shift toward preferential lipid storage in LGMD R2 (30, 31, 34), highlighting a major role of lipid metabolism in LGMD R2 pathophysiology. While the cause of these dysregulations is still unknown, our discovery that the major clathrin adaptor responsible for cholesterol uptake is increased in LGMD R2 muscle suggests the possibility that modulation of cholesterol levels could prove to be a possible therapeutic intervention. Although further investigations are needed to explore the functional consequences of DAB2 modulation on cholesterol homeostasis or lipid accumulation in LGMD R2, our study proposes DAB2 as a promising target for the development of novel therapeutic strategies.

In conclusion, our findings suggest that DAB2 could serve as a marker of disease progression. Overall, these results demonstrate that DAB2 is inversely correlated to dysferlin levels, indicating its potential involvement in the pathophysiology of LGMD R2. From this observation and the known functions of DAB2, we hypothesize that the increased expression of DAB2 may play a crucial role in impaired cholesterol homeostasis in LGMD R2 patients by enhancing cholesterol uptake via the LDLR endocytosis. If confirmed, these findings could represent a significant breakthrough for LGMD R2 patients and open the door to a novel therapeutic approach that may positively influence disease progression and reduce the associated burden.

## METHODS

### Sex as a biological variable

Our study examined male mice because male animals exhibited less variability in phenotype. Sex was not considered as a biological variable for patient muscle biopsies.

### Cell lines and culture

Experiments were performed using hiPSC lines from three unaffected individuals ("controls", referenced as WT1, WT2, WT3), and three LGMD R2 patients (referenced as KO_1, KO_2, MIS). WT1 and WT2 hiPSC lines were reprogrammed from healthy IMR-90 lung fibroblasts obtained from the ATCC Cell Lines Biology Collection (Washington, USA) and from GM1869 provided by the Coriell Cell Repository (Camden, USA)(63, 64). The WT3 hiPSC line is a commercial cell line supplied by Phenocell (Grasse, France). The hiPSC LGMD R2 KO_1 (heterozygous nonsense c.5713C>T, p.Arg1905X; c.3517dupT, p.Ser1173PhefsX2) and KO_2 (heterozygous nonsense c.5946 +1G>A; c.5497G>T; p.Glu1833X) are among the dysferlin-deficient lines deposited by the Jain Foundation with the WiCell Research Institute (Madison, USA), officially designated as Jain Foundation line JFNY1 (CDI#01456.103.11) and JFRBi1 (CDI#01457.101.08), respectively. The hiPSC LGMD R2 MIS line (homozygous missense c.4022T>C, p.L1341P) was provided by Dr. Simone Spuler (Max Delbruck Center, Berlin, Germany). Control and LGMD R2 hiPSC lines were maintained and expanded as single cells on feeder-free, vitronectin-coated dishes (Gibco, USA) in StemMACS iPS-Brew XF medium (Miltenyi Biotech, Germany).

hiPSC were differentiated into skeletal muscle cells (skMC) using commercial media (Geneabiocell®) as described previously (45). Briefly, hiPSC were dissociated with StemPro Accutase (Gibco, USA), seeded on collagen I-coated plates (Biocoat, DB Biosciences, USA), and maintained for 10 days in skeletal muscle induction medium (SKM01, AMSBIO, United Kingdom) with a passage at day 7. Myogenic precursors were then dissociated with 0.05% trypsin (ThermoFisher Scientific, USA) and reseeded on collagen I-coated plates for 7 days in skeletal myoblast medium (SKM02, AMSBIO, United Kingdom) until freezing. For terminal differentiation, myoblasts were thawed on collagen I-coated glass slides in skeletal myoblast medium and, at confluence, incubated with skeletal muscle differentiation medium (SKM03, AMSBIO, United Kingdom) for additional 5-7 days. hiPSC, myoblasts (MyoB), and myotubes (myoT) were analyzed at days 0, 17, and 24 of differentiation, respectively. Maturation levels of myoblasts and myotubes were compared with a primary control muscle cell line (C25CL48 myoblasts) established from a human muscle biopsy and provided by the MyoLine immortalization platform of the Institute de Myology (Paris, France), with informed consent and anonymization prior to immortalization, in accordance with EU GDPR.

To evaluate lipid metabolism, hiPSC-derived myotubes at day 3 were cultured in fatty-acid-supplemented SKM03 for 48 hours following a standardized protocol. Linoleic acid (LA, C18:2) and oleic acid (OA, C18;1) were dissolved in ethanol to obtain 100mM stock solutions and then diluted in serum-free culture medium, yielding final fatty acid concentrations of 50µM. Control cells received medium containing an equivalent volume of ethanol.

### Human patient muscle samples

Skeletal muscle biopsies from 2 healthy control and 14 dysferlin-deficient patients were obtained from the Unité de Morphologie Neuromusculaire, Institut de Myologie (Paris, France). Muscle biopsies from patients followed at Pitié-Salpêtrière Hospital were selected based on: (1) confirmed biallelic *DYSF* pathogenic variants, (2) absent or nearly absent dysferlin by immune histochemistry and/or western blot, (3) clinical evidence of neuromuscular disease across a broad range of ages, age at onset, and clinical severity. Open biopsies were obtained from the deltoid muscle, snap-frozen in isopentane cooled in liquid nitrogen, and used for histoenzymology and immunohistochemistry.

### Animal models

B6.A-Dysfprmd/GeneJ (strain 012767; referred to as Bla/J) and C57BL/6J (WT/control) male mice were housed in an SPF barrier facility with a 12-hour light/ dark cycle at the Center for Exploration and Experimental Functional Research (CERFE, GIP GENOPOLE, France) and were provided with food and water ad libitum. All animals were handled according to French and European guidelines for human care and use of experimental animals (APAFIS#35896). For *in vivo* studies aimed at restoring full-length dysferlin expression by gene therapy, 1-month-old Bla/J mice received an equimolar mixture of AAV8_C5.12_HR5-hDysf and AAV8_HR3-DysfpA.SV40 vectors in saline at 1.10e^14^ vg/kg (viral genomes per kilogram) or saline alone (controls)via a single intravenous injection into the tail vein. Mice were euthanized 1 or 6 months after injection by cervical elongation. Tibialis Anterior (TA), psoas (PSO) and gluteus (GLU) muscles were collected. Left muscles were mounted transversely on a cork stopper by 6% tracaganth gum (Sigma, USA), frozen in isopentane cooled in liquid nitrogen for histology, sectioned at 8 μm on a cryostat (Leica), placed on slides, and stored at -80°C. Right muscles were frozen directly in liquid nitrogen in 1.5ml tubes for RNA and protein extraction. All samples were stored at -80°C.

### Flow Cytometry

Single-cell suspensions of hiPSC were prepared after chemical dissociation with accutase (Invitrogen, USA), centrifuged at 900 rpm for 5 minutes, and resuspended in 2% FBS (Sigma, USA) in cold PBS. Cells were stained with fluorescent dye-conjugated antibodies (<u>Table S2</u>) for 30 minutes on ice in the dark. Cells were washed in cold PBS, and analyzed on a MACSquant analyzer (Miltenyi Biotec, Germany). Data were analyzed with FlowJo Software (BD Biosciences, USA).

### Transient transfection

hiPSC-derived myotubes were transfected at day 3 by using Lipofectamine RNAiMAX (Invitrogen, USA) according to the manufacturer’s instructions, with siRNAs targeting DAB2 (ON-TARGETplus Human DAB2, Dharmacon, USA) or a nonspecific scrambled control (Invitrogen, USA). 48 hours after transfection, myotubes were frozen for western blotting or fixed for immunostaining.

### Quantitative PCR

RNA was extracted according to two different methods depending on sample type. Total RNA from *in vitro* cells was isolated using the RNeasy Mini kit (Qiagen, Germany) with on-column DNase I digestion. Total RNA from mouse muscle biopsies was extracted using the NucleoZOL (Macherey-Nagel, Germany) folowing the manufacturer’s instructions. Briefly, nitrogen-frozen muscle tissue was divided, one piece was lysed in NucleoZOL and homogenized using a FastPrep-24 instrument (MP Biomedicals, USA). The lysate was supplemented with water, incubated for 10 minutes at room temperature, and centrifuged for 15 minutes at 12,000g. The upper phase was collected, incubated with isopropanol for 10 minutes at room temperature, and centrifuged 10 minutes at 12,000g to precipitate RNA. The pellet was washed with 75% ethanol, centrifuged 3 minutes at 8,000g, air-dried for 15 minutes, andresuspended in RNase-free water.

RNA quantity and purity were assessed using a NanoDrop spectrophotometer. A total of 500 ng RNA was reversetranscribed using SuperScript III (Invitrogen, USA). qPCR was performed on a QuantStudio 12K Flex (Applied Biosystems, USA) using Luminaris Color HiGreen qPCR Master Mix (Thermo Scientific, USA) or TaqMan gene expression Master Mix (Roche, Switzerland), following manufacturers’ instructions. Gene expression was quantified by the ΔCt method and normalized to human *18S* or murine *PO* expression. Primer sequences designed for this study are listed in <u>Table S3</u>. Sequences of other genes are commercially available (Applied biosystem, USA): *DYSF* (Hs01002513), *TTN* (Hs00399225) and *18S* (assay HS_099999).

### Immunostaining assay

Skeletal muscle cells were fixed with 4% paraformaldehyde (Euromedex, France) for 10 minutes at room temperature. After 3 washes in phosphate-buffered saline (PBS), cells were permeabilized with 0.5% Triton X-100 (Sigma, USA) for 10 minutes and blocked in PBS supplemented with 1% bovine serum albumin (BSA, Sigma, USA) for 1 hour at room temperature. Cells were stained for specific markers overnight at 4°C using primary antibodies (<u>Table S2</u>). After three washes in PBS, labeling was revealed with appropriate fluorophore-conjugated secondary antibodies (<u>Table S2</u>) for 1 hour at room temperature in the dark, and nuclei were counterstained with Hoechst (Invitrogen, USA).

Imaging was performed on a CellInsight CX7 imager (Cellomics Inc) with 20x or 40x objectives. For high-resolution acquisitions, coverslips were mounted in Fluoromount (Thermo Scientific, USA) and imaged on an LSM 880 confocal microscope (Zeiss, Germany) with a 63x oil-immersion objective and Z-stack acquisition. Z-stacks were combined using Zen Black software (Zeiss, Germany) to generate a single resolved image or for 3D myotube modeling.

### Lipid droplet analysis

After 48 hours of fatty acid supplementation, myotubes were immunostained with anti-desmin antibody, as described. During secondary antibody incubation, lipid droplets were labeled with LipidSpot™ 488 (Biotium, USA) for 1 hour at room temperature. Images were acquired on an LSM 880 confocal microscope (Zeiss, Germany) with a 63x oil immersion objective and Z-stack module. Z-stacks were automatically reconstructed in 3D using the 3D viewer module of Imaris analysis software (v.10.2, Oxford instruments, United Kingdom). Nuclei (Hoechst) were segmented using the *Add New Surfaces* function, adjusting thresholding parameters to optimize signal detection. Myotubes (desmin staining) were segmented similarly, with threshold adjustment and removal artifacts to generate a region of interest (ROI). Lipid droplets (LipidSpot) were analyzed using two complementary approaches. First, droplets were detected as individual puncta with the *Spots* module using optimized detection parameters (estimated spot diameter, background subtraction, threshold, quality filter). Second, droplets were segmented as continuous structures with the *Add New Surfaces* module using intensity thresholding. Lipid droplets were quantified within the ROI “myotubes” by using the *shortest distance to surface* filter. Detection parameters were applied uniformly across the image batch. Quantitative metrics (nuclei count, total myotube volume, number of lipid droplet spots and lipid volume within myotubes using the “Shortest Distance to Surface” filter) were extracted from the *Statistics* tab and exported for analysis.

### Histological characterization and immunohistochemistry

For hematoxylin-phloxine- saffron (HPS) staining, slides were thawed, air-dried for 10 minutes, and incubated for 5 minutes in Harris hematoxylin bath (Sigma, USA). Sections were washed in distilled water for 2 minutes, dipped in 0.2% (v/v) hydrochloric alcohol for 10 seconds, and rinsed in water for 1 minute. Slides were immersed in Scott’s water (0.5 g/L sodium bicarbonate and 20 g/L magnesium sulfate) for 1 minute, rinsed in water for 1 minute, stained in 1% (m/v) phloxine (Sigma, USA) for 30 seconds. Sections were rinsed in water for 1 minute 30 seconds, dehydrated in 70% ethanol for 1 minute and then absolute ethanol for 30 seconds. Finally, staining was completed in 1% saffron (v/v in absolute ethanol) for 3 minutes, followed by a wash in absolute ethanol. Sections were cleared in xylene for 2 minutes and mounted in Eukitt medium (Labonord, France). Slides were dried ≥24 hours and scanned using an AxioScan (Zeiss, Germany).

For immunohistochemical detection of dysferlin and DAB2, 10-µm transverse sections of human muscle biopsies were rehydrated with PBS for 10 minutes at room temperature, fixed with 4% paraformaldehyde (Euromedex, France) for 10 minutes, and immunostained as described. Coverslips were mounted in Fluoromount s(Thermo Scientific, USA) and images were acquired using an LSM 880 confocal microscope (Zeiss, Germany) with a 20x objective.

### Oil Red O staining on frozen muscle sections

Frozen human and mice muscle biopsy sections were air-dried for 15 minutes at room temperature and subsequently fixed in 60% isopropanol for 1 minute. Sections were then incubated in freshly prepared ORO working solution for 10 minutes, followed by a 30-second rinse in 60% isopropanol and a gentle wash in distilled water. Counterstaining was performed with hematoxylin for 30 seconds, after which sections were rinsed again in distilled water. Slides were mounted with Fluoromount-G, applied carefully to avoid displacement of the section or introduction of air bubbles, and coverslipped without applying pressure. Slides were dried under a chemical hood before imaging using an Evos cell imaging system (Thermo Fisher Scientific, USA).

### Western blot analysis

Proteins were extracted using two methods depending on sample origin. For whole-cell lysates of skeletal muscle, proteins were extracted with NP40 lysis buffer (Thermo Scientific, USA) supplemented with protease inhibitors (Complete PIC, Roche, Switzerland). Protein concentration was determined using Pierce BCA Protein Assay Kit (Thermo Scientific, USA) and absorbance measured at 562 nm on a CLARIOstar® microplate reader (BMG Labtech, Germany). A total of 20 μg of protein was separated using 4%–15% Criterion™ XT Tris-Glycine gels (BioRad, USA) and then transferred to PVDF membranes (BioRad, USA) using a Trans-Blot Turbo system (BioRad, USA). For mouse muscle biopsies, 30-µm sections were lysed in RIPA buffer (ThermoFisher, USA) supplemented with protease inhibitors (Complete PIC EDTA-free; Roche, Switzerland) and Benzonase (Merck-Millipore, Germany), then homogenized with a FisherBrand instrument (ThermoFisher, USA). Protein concentration was determined by BCA assay (Thermo Scientific, USA) and measured on an EnSpire multimode plate reader (PerkinElmer, USA). A total of 15 μg of protein was separated on 4%–12% NuPAGE Novex Tris-Bis gels (ThermoFisher, USA) and transferred to nitrocellulose membranes (ThermoFisher, USA) using an iBlot device (ThermoFisher, USA). Membranes were blocked in Odyssey blocking buffer (LI-COR, USA) for 1 hour at room temperature, incubated with primary antibodies (<u>Table S2</u>) diluted in blocking buffer overnight at 4°C or for 2 hours at room temperature. Washing was carried out 3 times for 10 minutes in TBS + 0.1% Tween-20 (VWR, USA) and membranes were incubated with appropriate fluorescent secondary antibodies (<u>Table S2</u>) for 1 hour at room temperature. After washing, proteins were detected by fluorescence (Odyssey CLx, LI-COR, USA).

### Transcriptomic analysis

Differential gene expression between hiPSC-derived myotubes was assessed using QuantSeq 3’mRNA sequencing on an Ion Torrent platform. For each of 18 samples, 100 ng total RNA was used to prepare libraries with the QuantSeq 3’ mRNA-Seq Library Prep Kit for Ion Torrent, generatingNGS libraries from the 3’ end of polyadenylated RNA. Libraries were amplified and barcoded (13 cycles) and quantified with the Agilent High Sensitivity DNA kit. Ten libraries (100 pM each) were pooled; emulsion PCR and enrichment were performed on the Ion OT2 instrument using the Ion PI Hi-Q OT2 200 kit (Thermo Fisher Scientific, USA). Samples were loaded on an Ion PI v3 Chip and sequenced on an Ion Proton System using Ion PI Hi-Q 200-bp chemistry (Thermo Fisher, USA). Quality control was performed with FastQC (v0.11.2). Reads were trimmed with Prinseq (v0.20.4; --trim-right 20) and filtered by Phred score (--trim-qual 20)(65). Reads were aligned to the Ensembl GRCh37.87 reference (protein-coding genes only: 19,311 genes; 143,832 transcripts) using RNA-STAR (v2.4.1d)(66) and filtered with samtools (v0.1.19)(67). Gene-level counts were obtained with HTSeq-count (v0.8.0)(68). Transcript expression was quantified with Kallisto (v0.43.1)(69). Finally, differential gene and transcript expression (DEG) between conditions was analyzed with DESeq2 (v1.18.1 using R v3.4.1)(70). Genes were considered significantly differentially expressed at adjusted P < 0.05 and |log2 fold change| > 0.4 (lfcThreshold = 0.4; altHypothesis = "greaterAbs"). Quantseq data generated in this study have been deposited on NCBI GEO under the accession number no.GSE261832.

Biological interpretation of differentially expressed genes was conducted with EnrichR (71) usingthe KEGG2021 database. Gene enrichment of up- and down-regulated DEGs was also performed using the over-representation approach in the clusterProfiler package (72), focusing on Gene Ontology (GO) Biological Process terms. GO terms with adjusted P < 0.1 were considered significant.

### Cross-referencing transcriptomic data

One previously published dataset (47, 48) was analyzed. This set containes Affymetrix mRNA profiles from 117 patient muscle biopsies using HG-U133A and HG-U1333B microarrays (n = 234 microarrays total; GEO accession no. GSE3307). While the original study analyzed 13 groups, here we analyzed only healthy and LGMD R2 skeletal muscle samples (Healthy: n=13, LGMD R2: n=10). Differential expression analysis (LGMD R2 vs. WT) was performed using GEO2R (LIMMA) on the GEO platform.

### Statistics

Data are presented as mean ± SD. Statistical analyses were performed using the Mann-Whitney test or an unpaired t-test with Welch’s correction. For comparisons involving more than two groups, one-way ANOVA was used followed by Dunnett’s multiple comparisons test. Data were log-transformed when necessary. Differences were considered significant at P-values of * *p* ≤ 0.05, \*\**p* ≤ 0.01, \*\*\**p* ≤ 0.001. “NS” indicates not significant. Correlations between data were assessed using Pearson correlation with significance set at \**p* ≤ 0.05. All graphs were generated in GraphPad Prism (v9.2.0).

### Study approval

All human cell lines were obtained from commercial collections as described or from collaborators with the subjects’ agreement, as evidenced by their signature of an informed consent form, and anonymised in accordance with the EU GDPR regulation. Human skeletal muscle biopsies were obtained from the Unité de Morphologie Neuromusculaire (Institut de Myologie, Paris, France). Biopsies were performed as part of the diagnostic process, and patients provided informed consent for genetic analysis and the reuse of biological samples for research purposes. All animals were handled in accordance with French and European guidelines for the care and use of experimental animals (APAFIS#35896).

## Supporting information

Movie S2_3D visualization of DAB2 staining in LGMD R2 KO myotubes

Movie S2_3D visualization of DAB2 staining in WT myotubes

Movie S5_3D visualization of lipid droplet accumulation in LGMD R2 KO myotubes treated with siDAB2

Movie S5_3D visualization of lipid droplet accumulation in WT myotubes

Movie S5_3D visualization of lipid droplet accumulation in LGMD R2 KO myotubes

## Data availability

All data generated or analyzed are included in this article or its supplemental material, including the Supporting Data Values file. The transcriptomic dataset generated and analyzed during this study is available in the NCBI’s Gene Expression Omnibus repository (GSE261832).

## Author contributions

C.B. and X.N. were responsible for the experimental conception and design. X.N. and I.R. were responsible for project management. C.B. performed the cell culture experiments, generated the cell banks, developed the pathological skeletal muscle model and realized the functional analyses. M.J. technically carried out the transcriptomic analysis. M.J., H.P., C.B., Q.M. and A.B. performed analysis and interpretation. E.P., M.B., N.G. provided technical assistance for cell culture. I.R. provided Bla/J mouse model, N.B. and V.A. conceived and performed animal experiments. C.B. and N.B.. investigated the mouse biopsies and carried out molecular analyses. S.V. has contributed his expertise on clathrin-mediated endocytosis linked to DAB2. T.E., G.F. and T.S. provided human muscular biopsies. C.B. conceived and performed the human biopsy analyses. C.B. prepared the figures. C.B. and X.N. wrote the manuscript. All authors reviewed and edited the paper.

## Acknowledgments

The authors thank Marc Bartoli (Marseille Medical Genetics, Marseille) for helpful discussions, Lina El Kassar, Jerome Polentes, Olivier Chose, Celine Leteur and Karine Giraud-Triboult from the research and innovation teams of I-Stem for providing expertise and technical support in imaging, quality controls and bioproduction, the Platform for Immortalization of Human Cells from the Centre of Research in Myology (Institute of Myology, Paris) for providing immortalized myoblasts. We thank Simone Spuler for providing some of the cellular models used in this study. We thank the Jain Foundation and Cellular Dynamics for generation, characterization, and provision of JFNY1 and JFRBi1 hiPSC. We also thank France Leturcq for providing help for mRNA extraction in patient muscle biopsies. We are grateful to the “*in vivo* department” of Genethon for technical support in animal experiments and to the AAV production department for viral productions. Some figures have been created with the BioRender website.

