## Supplemental file for "DAB2 as a biomarker and mechanistic link between lipid dysregulation and disease progression in LGMD R2"

**This PDF file includes:**

Figure S1. LGMD R2 hiPSC lines maintain pluripotency
Figure S2. Efficient skeletal myogenic differentiation of LGMD R2 hiPSC lines.
Figure S3. Transcriptomic profiling reveals differential gene expression in LGMD R2 myotubes.
Figure S4. DAB2 transcript is upregulated in LGMD R2 patient muscle biopsies.
Figure S5. DAB2 expression correlates with patient involvement.
Figure S6. Histological characterization of dysferlin-deficient Bla/J mice.
Figure S7. Dysferlin AAV gene therapy restores DAB2 in Bla/J mice.
Figure S8. Lipid accumulation occurs in the most affected muscles of Bla/J mice.
Figure S9. Dysferlin AAV gene therapy normalizes lipid accumulation in Bla/J mice.
Figure S10. siDAB2 treatment normalizes lipid accumulation in LGMD R2 myotubes.

Table S1. Clinical characteristics of patients with dysferlin deficiency.
Table S2. List of antibodies.
Table S3. List of primers.

### Supplementary Figure 1

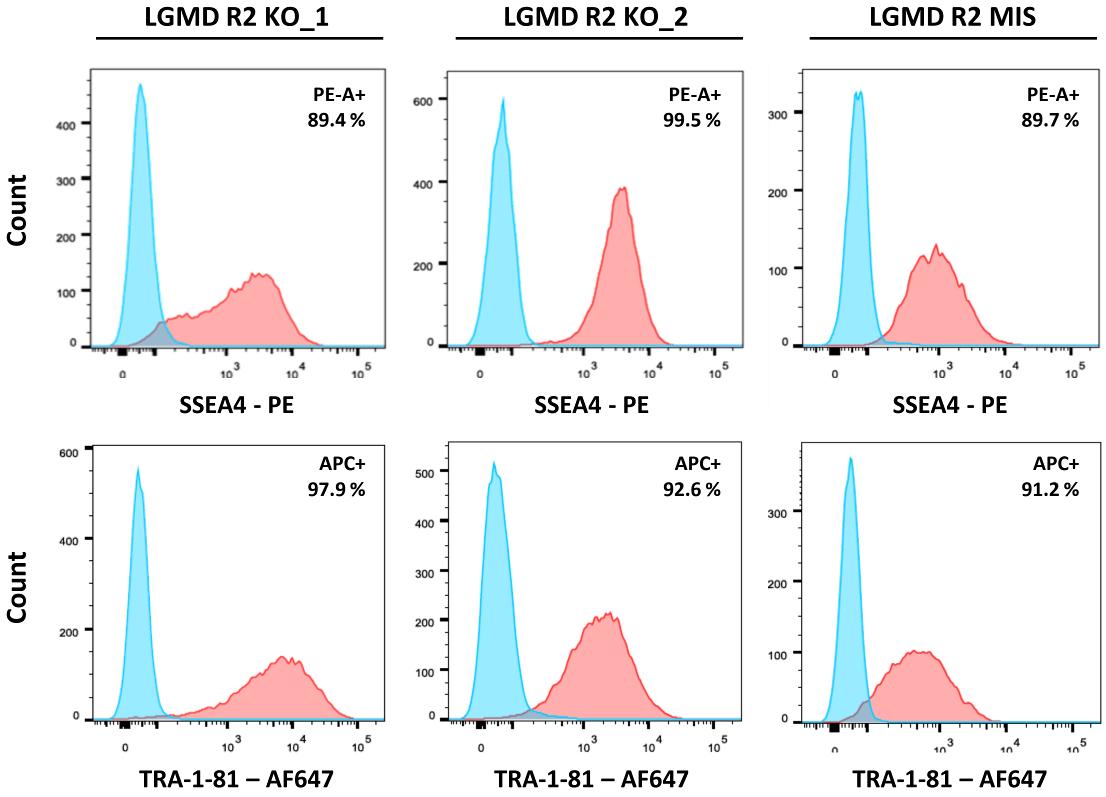


**Figure S1. LGMD R2 hiPSC lines maintain pluripotency.** Flow cytometry analysis of surface pluripotency markers SSEA4 (PE-A+) and TRA-1-81 (APC+) in three LGMD R2 hiPSC lines. Blue histograms represent isotype controls, and red histograms represent experimental samples.

### Supplementary Figure 2

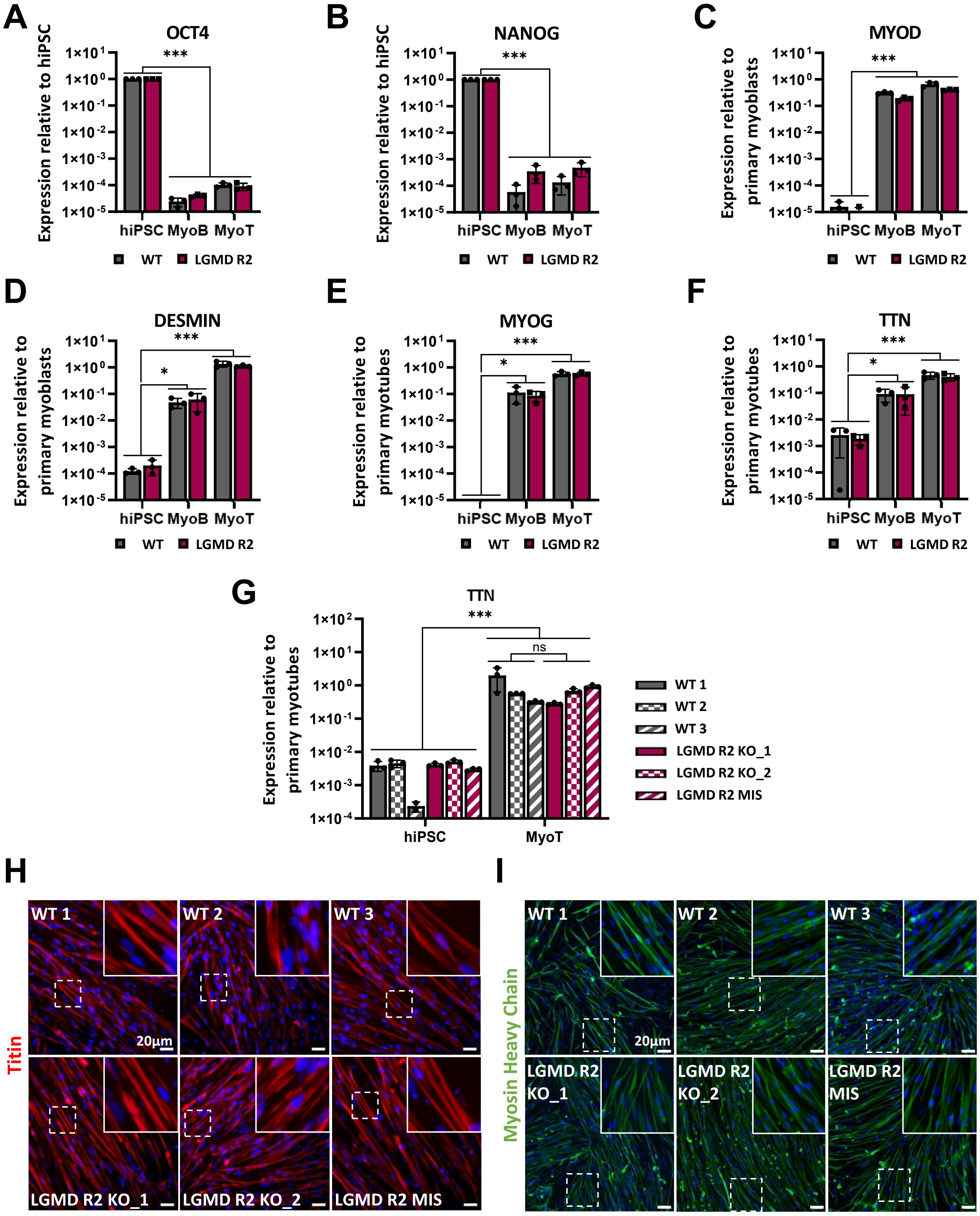

**Figure S2. Efficient skeletal myogenic differentiation of LGMD R2 hiPSC lines.** (**A-F**) mRNA expression levels of pluripotency marker OCT4 (**A**) and NANOG (**B**), and myogenic markers MYOD (**C**), DESMIN (**D**), MYOG (**E**), and TTN (**F**) at day 0 (hiPSC), day 17 (myoblasts), and day 24 (myotubes). Gene expression was normalized to hiPSC, primary myoblasts, or primary myotubes, respectively. Data are shown as mean ± SD of three control (grey) or three LGMD R2 (pink) lines (n=3). *p ≤ 0.05, ***p ≤ 0.001 (Nonparametric Mann-Whitney test). **(G)** TTN expression measured by qPCR at day 0 (hiPSC) and day 24 (myotubes), normalized to primary myotubes. Data represent mean ± SD of three skMC differentiations per cell line (n=3). *p ≤ 0.05 hiPSC vs. MyoT group (Nonparametric Mann-Whitney test); ns p > 0.05 WT vs. LGMD R2 group at myotube stage (Nonparametric Mann-Whitney test). Characterization of titin (red) (**H**) and myosin heavy chain (MHC) (**I**) expression by immunostaining in control and LGMD R2 hiPSC-derived myotubes (day 24). Nuclei were stained with Hoechst (blue). White boxes indicate magnified areas. Scale bar: 20 µm.

Abbreviations: hiPSC, human induced pluripotent stem cells; MyoB, myoblasts; MyoT, myotubes.

### Supplementary Figure 3

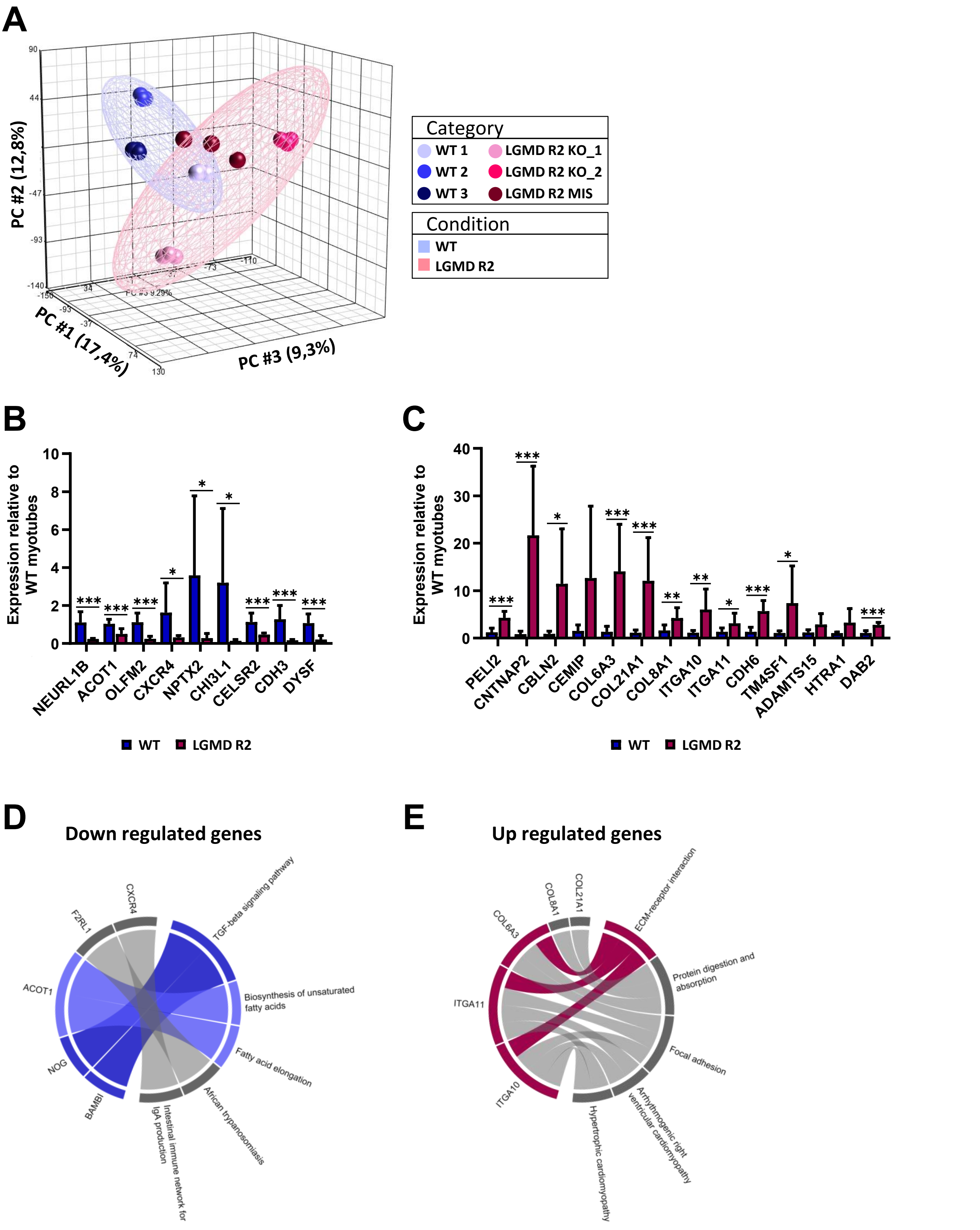

**Figure S3. Transcriptomic profiling reveals differential gene expression in LGMD R2 myotubes.** (**A**) Principal component analysis (PCA) of transcriptome profiles obtained from myotubes derived from three control (blue) and three LGMD R2 (pink) lines. Each point represents an independent skMC differentiation (n=3 per cell line). Samples with similar gene expression profiles cluster together. (**B-C**) Validation of 23 out of 52 differentially expressed genes by quantitative PCR. Genes down-regulated (**B**) or up-regulated (**C**) in LGMD R2 myotubes are shown. Gene expression was normalized to the mean of myotubes derived from three control lines. Data are represented as mean ± SD of three myogenic differentiations from three control lines (blue) or three LGMD R2 lines (pink) (n=9). $^{*}p \leq 0.05$, $^{**}p \leq 0.01$, $^{***}p \leq 0.001$ (Nonparametric Mann-Whitney test). (**D-E**) KEGG 2021 Human enrichment analysis of 13 significantly down-regulated genes (**D**) and 39 significantly up-regulated genes (**E**) in LGMD R2 myotubes. The top five enriched pathways ($p < 0.05$) are shown. Lines connect genes to the pathways in which they are involved. Pathways relevant to pathology are highlighted in blue (down-regulated) or pink (up-regulated).

### Supplementary Figure 4

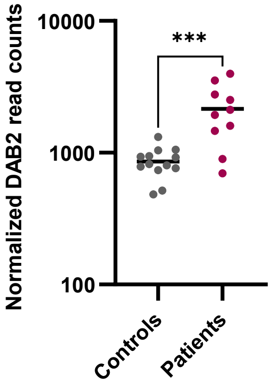


**Figure S4. DAB2 transcript is upregulated in LGMD R2 patient muscle biopsies.** Analysis of DAB2 expression (read counts) from a previously published Affymetrix transcriptomic dataset comparing 10 muscle biopsies from LGMD R2 patients (pink) and 13 unaffected individuals (grey). Each dot represents an individual biopsy. ***$p \leq 0.001$ (Unpaired t-test with Welch's correction).

### Supplementary Figure 5

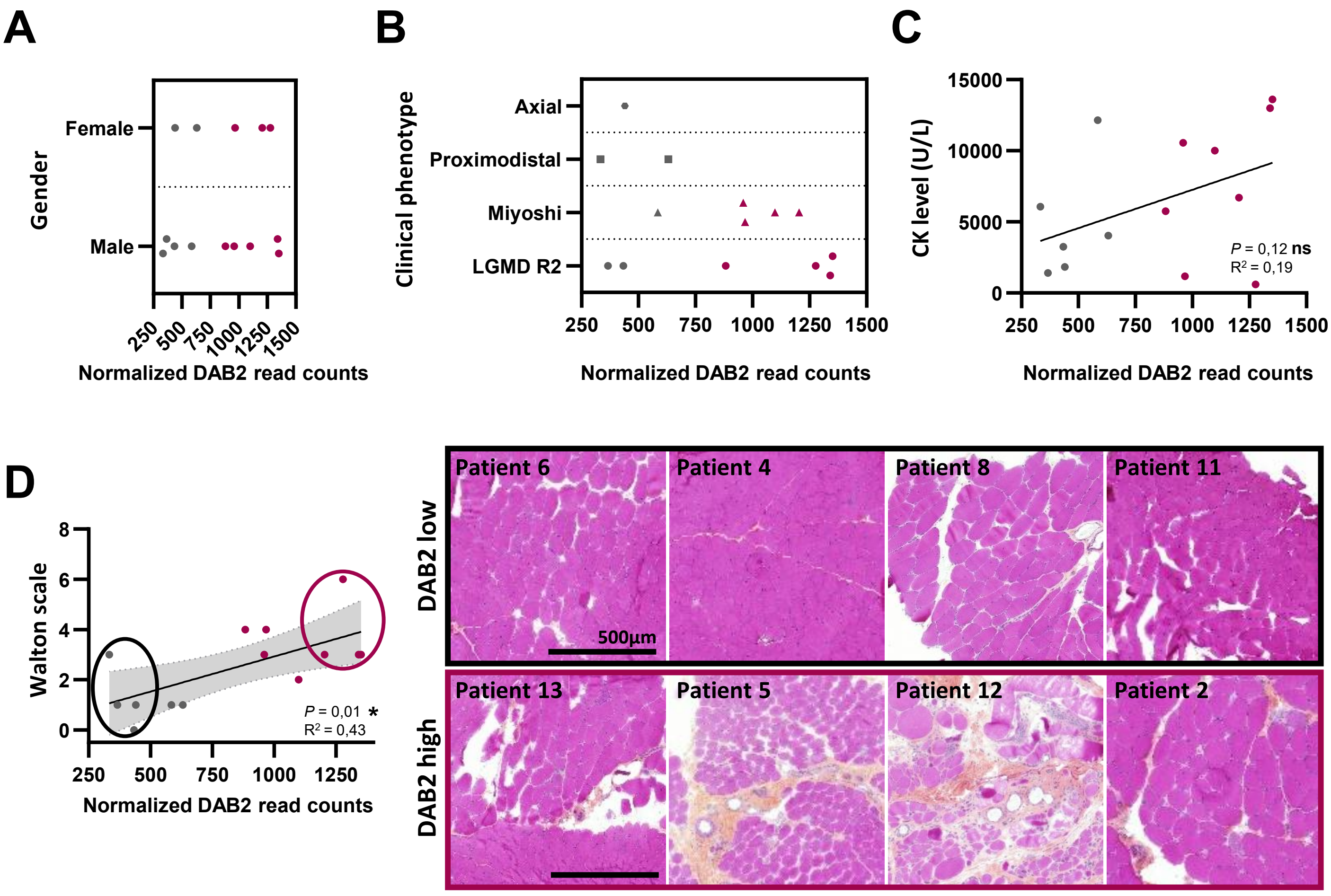


**Figure S5. DAB2 expression correlates with patient involvement.** Analysis of DAB2 expression in dysferlin-deficient muscle biopsies according to gender (**A**), clinical phenotype (**B**), or creatine phosphokinase levels (**C**). Each dot represents an individual biopsy. DAB2 expression is color-coded from low (grey) to high (pink). No significant correlations were observed (ns, $p > 0.05$, Pearson correlation analysis). (**D**) Visualization of DAB2 expression relative to patient involvement based on the Walton scale (poorly involved 0 ≤ Walton scale ≤ 10 severely involved). Representative HPS-stained muscle sections are shown for patients with the lowest (grey circle) and highest (pink circle) DAB2 expression. Scale bar = 500 µm. *$p \leq 0.05$ (Pearson correlation analysis). Abbreviation: CK, creatine kinase.

### Supplementary Figure 6

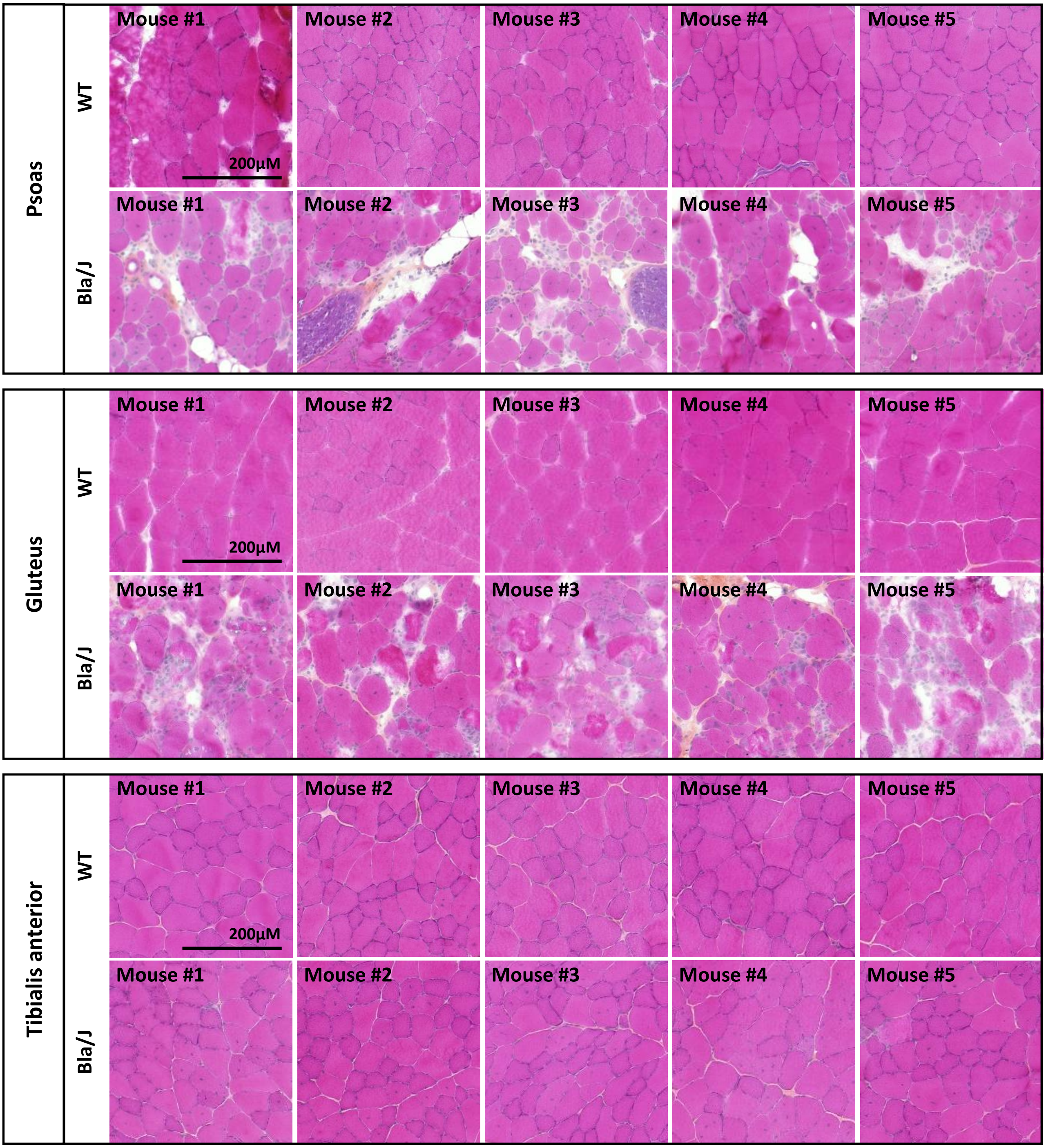


**Figure S6. Histological characterization of dysferlin-deficient Bla/J mice.** Comparative analysis of psoas, gluteus and tibialis anterior muscles from six-month-old control and Bla/J mice using HPS staining. Scale bar = 200 µm.

### Supplementary Figure 7

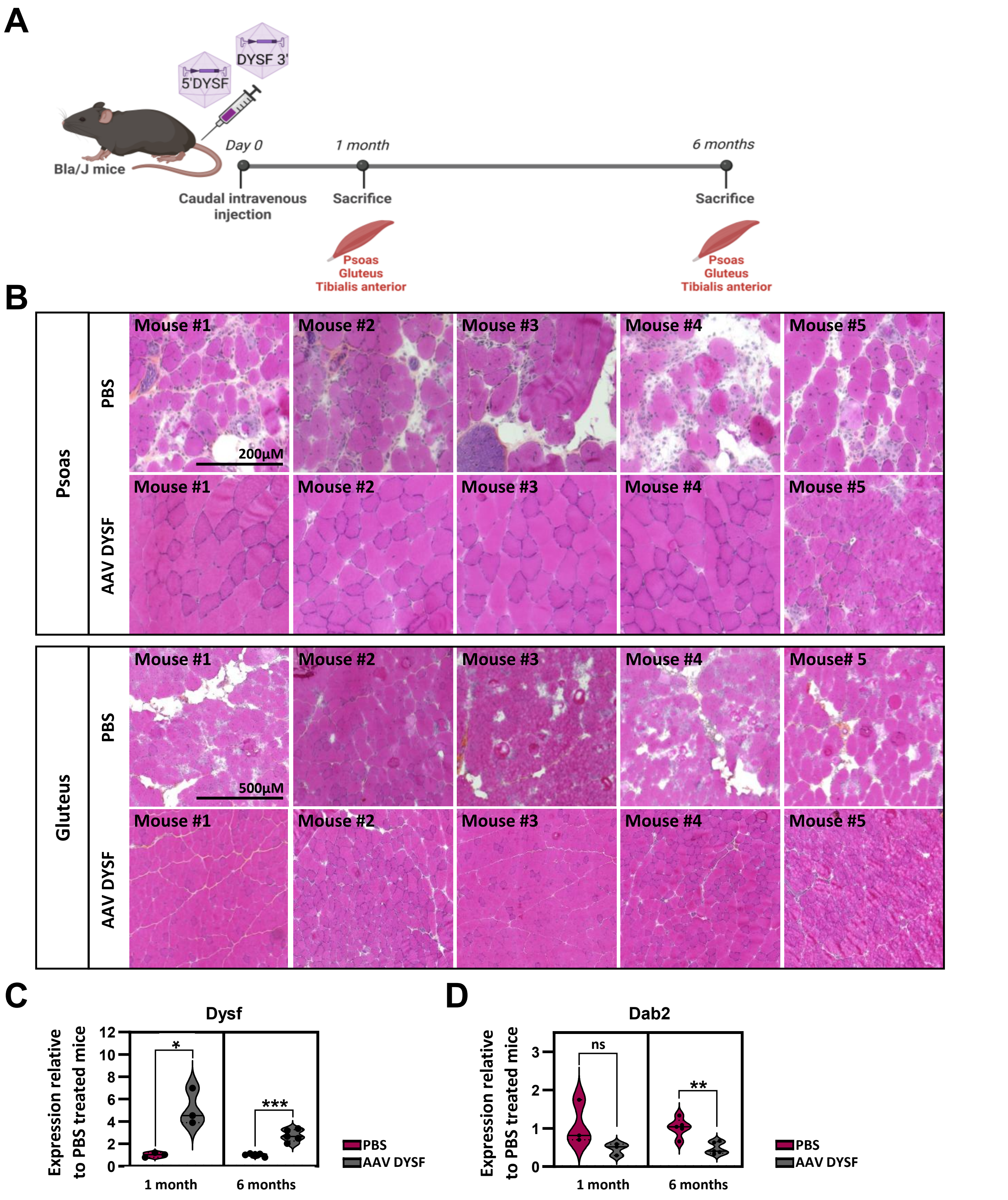

**Figure S7. Dysferlin AAV gene therapy restores DAB2 in Bla/J mice.** (**A**) Schematic representation of the gene therapy protocol for one-month-old Bla/J mice using a dual AAV vector to restore full-length dysferlin. Mice received a single treatment; protocol duration is indicated. Created with BioRender.com. (**B**) Representative HPS-stained sections of psoas and gluteus muscles from Bla/J mice six months after treatment with PBS or dual AAV dysferlin rescue (AAV DYSF). Scale bar = 200 or 500 µm. (**C–D**) qPCR analysis of Dysf (**C**) and Dab2 (**D**) expression in gluteus muscle after one or six months of treatment. Expression was normalized to PBS-treated mice. Data represent mean ± SD of mice per condition (n = 3–5). *p ≤ 0.05 or **p ≤ 0.01, ***p ≤ 0.001 (Unpaired t-test with Welch's correction).

### Supplementary Figure 8

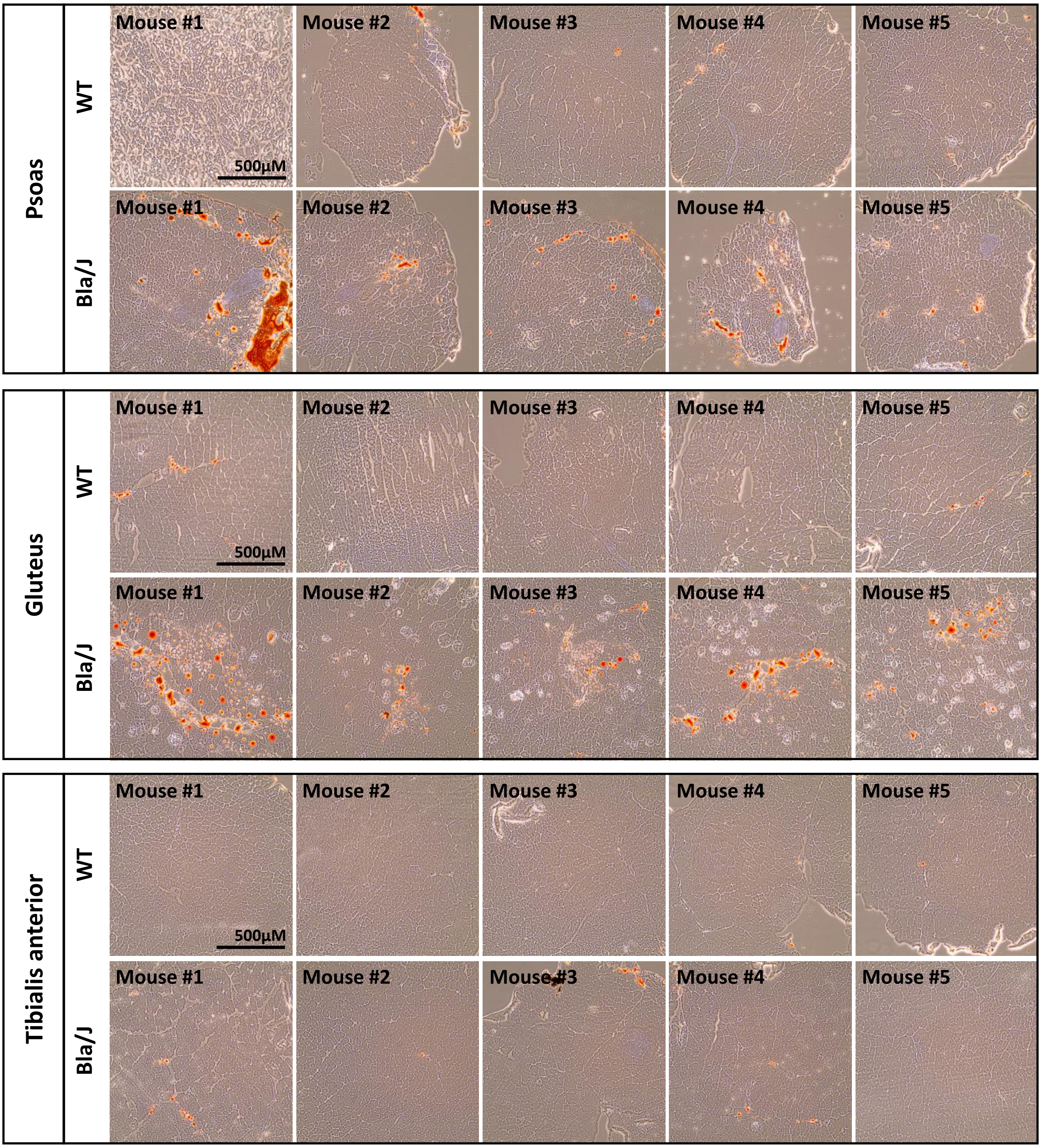


**Figure S8. Lipid accumulation occurs in the most affected muscles of Bla/J mice.** Representative Oil Red O-stained sections of psoas, gluteus, and tibialis anterior muscles from six-month-old WT and Bla/J mice. Scale bar = 500 µm.

### Supplementary Figure 9

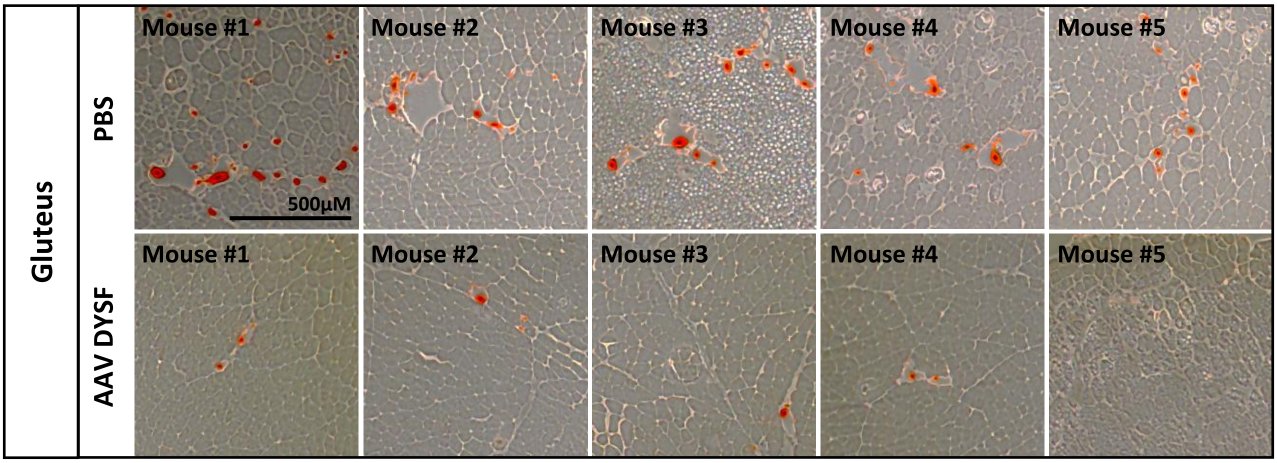


**Figure S9. Dysferlin AAV gene therapy normalizes lipid accumulation in Bla/J mice.** Representative Oil Red O-stained sections of gluteus muscles from Bla/J mice six months after treatment with PBS or dual AAV dysferlin rescue (AAV DYSF). Scale bar = 500 µm.

### Supplementary Figure 10

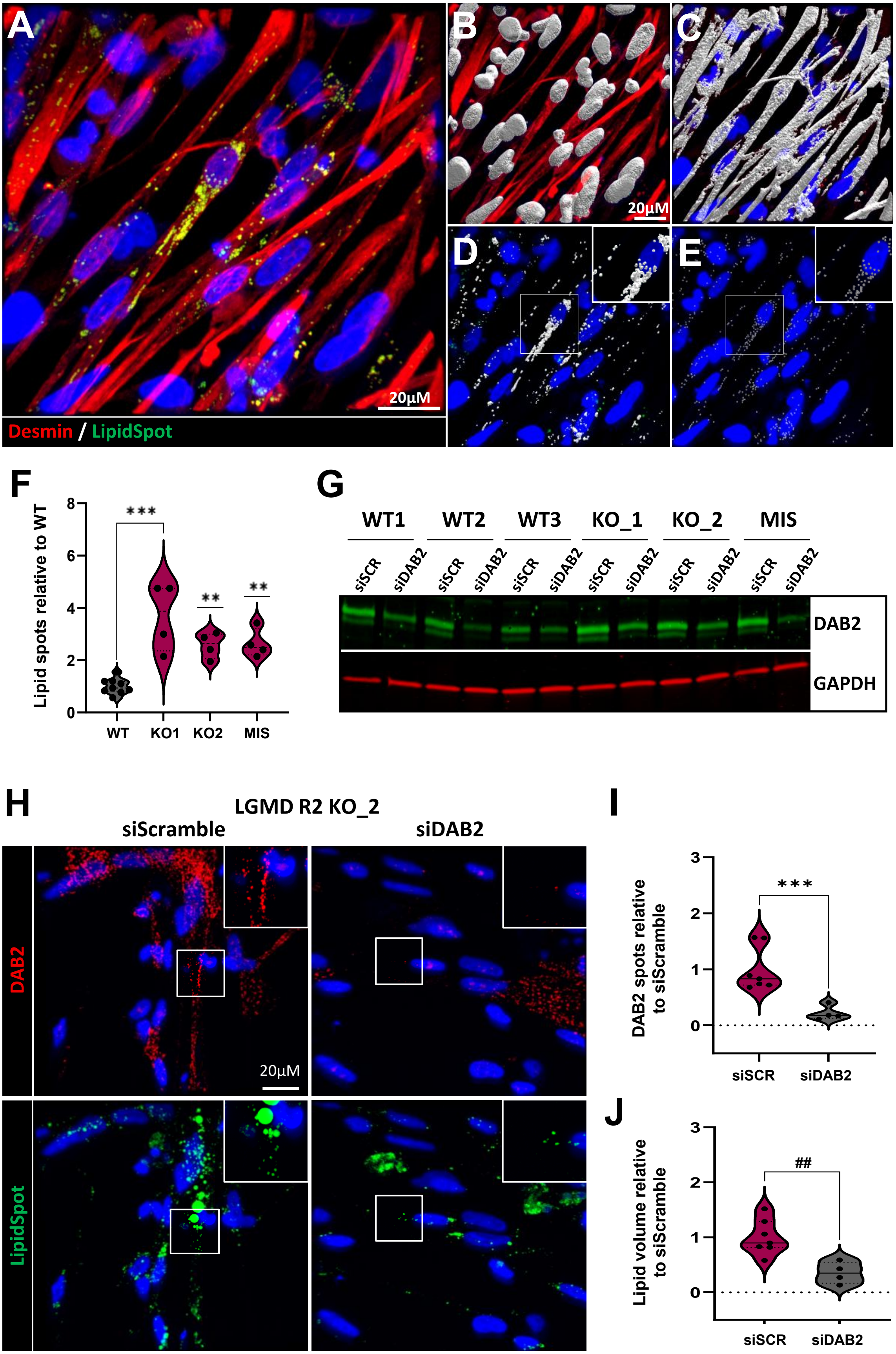

**Figure S10. siDAB2 treatment normalizes lipid accumulation in LGMD R2 myotubes.**

(**A-E**) Immunostaining of lipid droplets (LipidSpot, green) in hiPS-derived myotubes (desmin, red) supplemented with fatty acids for 48 hours (**A**). Nuclei were counterstained with Hoechst (blue). Imaris detection masks were used to identify nuclei (**B**), myotubes (**C**), and lipid droplets for quantification of lipid volume (**D**) and lipid droplet number (**E**). White boxes indicate magnified regions. Scale bar = 20 µm. (**F**) Quantification of lipid droplet number in WT and LGMD R2 hiPSC-derived myotubes. Data represent mean ± SD of a representative experiment from *n*=3 experiments. **p ≤ 0.01, ***p ≤ 0.001 (One-way ANOVA, Dunnett's multiple comparisons test). (**G-J**) Analysis of siDAB2 treatment on lipid accumulation in fatty acid-supplemented myotubes. (**G**) Representative immunoblot of human DAB2 P96 isoform expression (green) in healthy and LGMD R2 myotubes treated for 48 hours with siRNA targeting DAB2. GAPDH is a loading control. (**H**) Immunostaining of DAB2 (red) and lipid droplets (green) in LGMD R2 KO_2 hiPSC-derived myotubes after 48 hours of treatment with siScramble (left) or siDAB2 (right). White boxes indicate magnified regions. Scale bar = 20 µm. Associated quantification of DAB2 droplet number (**I**) and lipid volume (**J**) in LGMD R2 KO_2 hiPSC-derived myotubes after treatment with siScramble (pink) or siDAB2 (grey). Data represent mean ± SD of a representative experiment from *n*=3 experiments. ##p ≤ 0.01, ***p ≤ 0.001 (Unpaired t test; transformed data for lipid volume). Abbreviation: siSCR, siScramble.

### Supplementary Table 1

#### Table S1. Clinical characteristics of patients with dysferlin deficiency.

| Patient | Gender | Muscle biopsy | Clinical phenotype | Genetic mutation | Dysferlin expression (wb) | Age at disease onset (years) | Biopsy age (years) | Diagnostic at biopsy | CK level (IU/L) | Walton score |
|---|---|---|---|---|---|---|---|---|---|---|
| 1 | Male | Deltoide | LGMD R2 | HTZ: c.2998T>C / c.4795-1G>C | absent | 16 | 31 | Dystrophic | 5753 | IIII |
| 2 | Male | Deltoide | LGMD | HMZ: c.5341-2A>C | absent | 22 | 23 | Dystrophic | 13618 | III |
| 3 | Male | Deltoide | Miyoshi | HTZ: c.2643+1G>A / c.4577A>C | absent | 18 | 35 | Dystrophic | 10561 | III |
| 4 | Male | Deltoide | LGMD R2 | HTZ: c.6017G>A / Ø | absent | 3 | 15 | Aspecific myopathic | 1404 | I |
| 5 | Female | Finger expander | LGMD R2 | HMZ: c.6044T>C | absent | 17 | 36 | Dystrophic | 605 | VI |
| 6 | Male | Deltoide | Proximodistal | HMZ: del exon 55 | absent | 30 | 35 | Dystrophic | 6072 | III |
| 7 | Male | Anterior hamstring | Miyoshi myopathy | HTZ: c.3517dupT / c.5871_5872delGT | absent | 19 | 20 | Dystrophic | 12147 | I |
| 8 | Male | Deltoide | LGMD R2 | HTZ: c.757C>T / c.5813-5821del | trace | 35 | 33 | Aspecific myopathic | 3247 | 0 |
| 9 | Female | Deltoide | Miyoshi | HMZ: c.5314-5318del | absent | 37 | 56 | Aspecific myopathic | 1167 | IIII |
| 10 | Male | Deltoide | Miyoshi | HTZ: c.1911C>A / c.2641A>C | trace | 17 | 20 | Dystrophic | 10000 | II |
| 11 | Female | Deltoide | Axial | HTZ: c.906+4A>G / c.1813C>T | absent | 52 | 55 | Aspecific myopathic | 1828 | I |
| 12 | Male | Deltoide | LGMD R2 | HTZ: del exon 5 / del exon 5 | absent | 23 | 30 | Dystrophic | 13000 | III |
| 13 | Female | Deltoide | Miyoshi | HTZ: c.20T>C / c.2810C>T | trace | 18 | 24 | Dystrophic | 6710 | III |
| 14 | Female | Deltoide | Proximodistal | HTZ: del exon 5 / c.906+4A>G | absent | 37 | 41 | Dystrophic | 4043 | I |

| Patient | Gender | Muscle biopsy | Clinical phenotype | Genetic mutation | Dysferlin expression (wb) | Age at disease onset (years) | Biopsy age (years) | Diagnostic at biopsy | CK level (IU/L) | Walton score |
|---|---|---|---|---|---|---|---|---|---|---|
| 1 | Female | Deltoide | | | present | CONTROL | 25 | | | |
| 2 | Male | Deltoide | | | present | CONTROL | 32 | | | |

### Supplementary Table 2

**Table S2. List of antibodies.**

| Application | Antibodies | Supplier | Reference | Dilution |
|---|---|---|---|---|
| Flow cytometry | TRA-1-81 AF647 | Biolegend | 330706 | 1:25 |
| | AF647 Isotype control | Biolegend | 401618 | 1:250 |
| | SSEA4 PE | Miltenyi | 130-122-914 | 1:50 |
| | PE Isotype control | Miltenyi | 130-113-462 | 1:50 |
| Immunostaining | α-actinin | Sigma | A7811 | 1 :1 000 |
| | Dab2 | Abcam | ab33441 | 1 :200 |
| | Desmin | R&D systems | AF3844 | 1 :200 |
| | Dysferlin | Abcam | ab124684 | 1 :200 |
| | Dysferlin | Novocastra | HAMLET-CE | 1 :50 |
| | LipidSpot 488 | Biotium | 70065 | 1 :1000 |
| | Myosin heavy chain | DSHB | 3ea | 1 :50 |
| | Titin | US Biological | T5650 | 1 :100 |
| | Donkey anti-mouse AF488 | Invitrogen | A-21202 | 1 :1 000 |
| | Donkey anti-rabbit AF488 | Invitrogen | A-21206 | 1 :1 000 |
| | Donkey anti-mouse AF555 | Invitrogen | A-31570 | 1 :1 000 |
| | Donkey anti-rabbit AF555 | Invitrogen | A-31572 | 1 :1 000 |
| | Donkey anti-goat AF555 | Invitrogen | A-21432 | 1 :1 000 |
| Western Blot | Actin | Sigma | A2066 | 1 :300 |
| | Dysferlin | Novocastra | HAMLET-CE | 1 :500 |
| | Dab2 | Abcam | ab33441 | 1 :500 |
| | Dab2 | BD Biosciences | 610465 | 1 :100 |
| | GAPDH | Thermofisher | 39-8600 | 1 :500 |
| | IRDye® 680RD Donkey anti-Rabbit | Li-Cor | 926-68073 | 1 :10 000 |
| | IRDye® 800CW Donkey anti-Rabbit | Li-Cor | 926-32213 | 1 :5 000 |
| | IRDye® 680RD Donkey anti-Mouse | Li-Cor | 926-68072 | 1 :10 000 |
| | IRDye® 800CW Donkey anti-Mouse | Li-Cor | 926-32212 | 1 :5 000 |

### Supplementary Table 3

**Table S3. List of primers.**

**List of human primers.**

| Gene | Forward | Reverse |
|---|---|---|
| ACOT1 | ATCTGGAGTACTTTGAAGAAGCTGT | CCAACTCCTGGACCTTTTACCTCA |
| ADAMTS15 | AACAAGCACAGGGTGGATGG | TGCAGGATCGGTATTTCACCC |
| CBLN2 | ACCATCCAGGTCAGTTTAATGC | AACACCAAGAAGCCCGAGAA |
| CDH3 | CAGCTACCGCATCCTGAGAG | GTGGGAGGGCTTCCATTGTC |
| CDH6 | CCCGATTCCCCCAGAGTACAT | CGTCGCTGGCTTTGATTCTG |
| CELSR2 | ATGTGTGTGACCTGAACCCG | GGCTGGTCAATCCTGGTCTC |
| CEMIP | ACTCAGGGCTGTTGTTCCTG | TCGGTGAACTTGGGGTAAGC |
| CHI3L1 | AAGAACAGGAACCCCAACCTG | TGCTGTTTGTCTCTCCGTCC |
| CNTNAP2 | ACCCCAGATGGAAGCCCTTA | GATCTGTGCAGTTGCGTTCG |
| COL21A1 | CCTGGAACACCGGGATCTAA | TGCTCCTGGTTCTCCCTTGA |
| COL6A3 | TGCGGGAAGCAGGATAACAG | CATCATCCCCAGACCTGTCG |
| COL8A1 | CCTGGGTCAGCAAGTACCTC | TGGTATTTCTTTGCCTTTCTTGGG |
| CXCR4 | TGGTCTATGTTGGCGTCTGG | GTCATTGGGGTAGAAGCGGT |
| DAB2 | GACCAGATGGATTTGTTTGGGG | GAACAGGAGGTGATGCCGTT |
| DESMIN | ATTGGAGGACCGATTTGCC | TCACCGTCTTCTTGGTATGGA |
| HTRA1 | TGTCACTCACGTCCAGCAAA | GACATCATTGGCGGAGACCA |
| ITGA10 | GGGGGAGCAGATTGGTTCAT | GTTCCTTGGAGGGTCAGCAA |
| ITGA11 | AATAAGTGGCTGGTCGTGGG | GTGACCCTTCCCAGGTTGAG |
| MYOD | GGGGCTAGGTTCAGCTTTCT | CTACATTTGGGACCGGAGTG |
| MYOG | TAAGGTGTGTAAGAGGAAGTCG | CCACAGACACATCTTCCACTGT |
| NANOG | CAAAGGCAAACAACCCACTT | TCTGCTGGAGGCTGAGGTAT |
| NEURL1B | GTGAGTCCGTCCAGTTGAGG | GGGACTCGGATAAGACAGCG |
| NPTX2 | TTGGACAAGAGCAGGACACC | GAGCAGTTGGCGATGTTGAC |
| OCT4 | CCTCACTTCACTGCACTGTA | CAGGTTTTCTTTCCCTAGCT |
| OLFM2 | TGGAGCAGTACAAGGCAGAC | ATACCCGTAGGCACCCATCT |
| PELI2 | CGGATATGTTTCAGGTGGGC | CTGTGTGATCTGGGCTTCGT |
| TM4SF1 | GCAAACGATGTGCGATGCT | GGCCGAGGGAATCAAGACAT |
| 18S | TCTTCAGTCCGCTCCAGGTCT | GAGGATGAGGTGGAACGTGT |

**List of mouse primers.**

| Gene | Forward | Reverse |
|---|---|---|
| DYSF | GGTGTTCGAGAACCAGACCC | GCTGTACTCCCAGCCTTGTT |
| DAB2 | CTCAGGGACAACACAAGCAA | AAATTGATGCTGGCCTTCAC |
| PO | CTCCAAGCAGATGCAGCAGA | ATAGCCTTGCGCATCATGGT |
